# Deep sequencing of Phox2a nuclei reveals five classes of anterolateral system neurons

**DOI:** 10.1101/2023.08.20.553715

**Authors:** Andrew M. Bell, Charlotte Utting, Allen C. Dickie, Mateusz W. Kucharczyk, Raphaëlle Quillet, Maria Gutierrez-Mecinas, Aimi N.B. Razlan, Andrew H. Cooper, Yuxuan Lan, Junichi Hachisuka, Greg A. Weir, Kirsty Bannister, Masahiko Watanabe, Artur Kania, Mark A. Hoon, Iain C. Macaulay, Franziska Denk, Andrew J. Todd

## Abstract

The anterolateral system (ALS) is a major ascending pathway from the spinal cord that projects to multiple brain areas and underlies the perception of pain, itch and skin temperature. Despite its importance, our understanding of this system has been hampered by the considerable functional and molecular diversity of its constituent cells. Here we use fluorescence-activated cell sorting to isolate ALS neurons belonging to the Phox2a-lineage for single-nucleus RNA sequencing. We reveal five distinct clusters of ALS neurons (ALS1-5) and document their laminar distribution in the spinal cord using *in situ* hybridization. We identify 3 clusters of neurons located predominantly in laminae I-III of the dorsal horn (ALS1-3) and two clusters with cell bodies located in deeper laminae (ALS4 & ALS5). Our findings reveal the transcriptional logic that underlies ALS neuronal diversity in the adult mouse and uncover the molecular identity of two previously identified classes of projection neurons. We also show that these molecular signatures can be used to target groups of ALS neurons using retrograde viral tracing. Overall, our findings provide a valuable resource for studying somatosensory biology and targeting subclasses of ALS neurons.

**Significance Statement:** The anterolateral system (ALS) is a major ascending pathway from the spinal cord that underlies perception of pain, itch and skin temperature. It is therefore an important target for the development of new treatments for chronic pain. Our understanding of this system has been hampered by the considerable diversity of its constituent cells. Here we dissect the complex heterogeneity of these cells by using high-resolution RNA sequencing. We reveal five distinct types of ALS neurons, which are differentially distributed within the spinal cord, and probably represent functional populations. Our data provide novel insights into the molecular architecture of the ALS, and will be important for future studies to define the roles of different ALS cell types in sensory processing.

## Main Text Introduction

The anterolateral system (ALS) is a major pathway that transmits somatosensory information from the spinal cord to the brain and is crucial for pain, itch and the perception of skin temperature (1–3). Disruption of the ALS, both physically through anterolateral cordotomy (4) and chemically using targeted toxins (5), results in effective pain relief. Neurons of the ALS therefore represent an attractive target for novel therapeutics. These cells are located in various regions of the spinal cord, with the highest density in lamina I and the lateral spinal nucleus (LSN). Other ALS cells are scattered throughout the deep dorsal horn, ventral horn and the area around the central canal (1, 2, 6). Neurons of the ALS project extensively throughout the brain, where they engage nociceptive circuits (7). Their major targets include the thalamus, periaqueductal grey matter, lateral parabrachial area (LPb), and caudal ventrolateral medulla. Despite their importance as the main pain-related ascending output from the spinal cord, ALS neurons are vastly outnumbered by interneurons, since projection neurons only account for ∼1% of spinal neurons (8).

There have been many attempts to classify ALS neurons. Early studies examined a variety of parameters, including morphology, projection target and physiological properties. For example, electrophysiological recording from antidromically identified projection neurons in animals of various species, mostly under general anaesthesia, revealed a wide variety of response properties. The majority of these studies have emphasised convergence of multiple modalities (mechanical, thermal and pruritic), onto individual ALS neurons, although with marked differences in the responses of individual cells to different stimuli. However, there is also evidence that some cells have more restricted types of input, and may represent labelled lines. In particular, it has been suggested that while most lamina I ALS cells are activated by noxious thermal and/or mechanical stimuli (9, 10), some respond preferentially or exclusively to skin cooling (9, 11–14). There is also evidence that for ALS neurons in deep dorsal horn the response properties depend on the laminar location of dendritic arbors (15). One of the main points to emerge from these studies is the functional heterogeneity among ALS neurons.

More recently, attempts have been made to identify genetic markers that are selective for ALS neurons. Transcriptomic studies have revealed widespread expression of Lypd1 and Zfhx3 among these cells (16–18). In addition, it has been shown that within the spinal cord developmental expression of the transcription factor Phox2a is restricted to ALS neurons, and captures cells in both superficial and deep dorsal horn. This allows ALS neurons to be targeted through mouse genetics (19–21). Furthermore, recent studies have proposed molecular signatures for distinct functional populations within the ALS. These have led to the identification of potential marker genes, including *Tac1*, *Tacr1*, *Gpr83*, *Cck*, *Nptx2*, *Crh* and *Nmb* (22–24). Despite these advances, there is no unbiased, unifying, classification scheme that describes the organisational logic of the ALS. Here we used a transgenic approach to isolate Phox2a-dependent ALS neurons for single nucleus RNA sequencing (snRNAseq). Our results reveal 5 distinct populations, demonstrating fundamental transcriptional subdivisions within the ALS. These findings should advance our knowledge of how co-ordinated activity in this system leads to complex multidimensional percepts such as pain.

## Results

### Single nucleus sequencing of Phox2a-lineage neurons

In order to isolate and sequence Phox2a-derived ALS neurons, we first crossed the Phox2a::Cre mouse line with the Cre-dependent Sun1-GFP reporter mouse (Fig. 1A). This resulted in nuclear GFP labelling of Phox2a-lineage neurons in the dorsal horn with a laminar distribution pattern as expected from previous reports (Fig. S1A). Nuclei were isolated from 16 mice in total, 8 of which had undergone left-sided spared nerve injury (SNI), 8 days prior to tissue harvest. The SNI animals were included to assess whether substantial transcriptional changes could be identified within projection neurons in a neuropathic pain state. Equal numbers of male and female animals were included in each group. Spinal cords were removed and the entire left dorsal horn of the L4 segment was dissected rapidly and frozen on dry ice. Phox2a-GFP nuclei were isolated from the frozen spinal cord tissue using hypotonic cell lysis followed by fluorescence-activated cell sorting (FACS). We used DR staining to isolate nuclei from debris and then gated based on GFP fluorescence (Fig. S1B,C). Nuclei positive for GFP were rare, generally representing <0.2% of all nuclear events passing through the sorter, as expected from the low proportion of spinal neurons that belong to the ALS.

**Figure 1.**
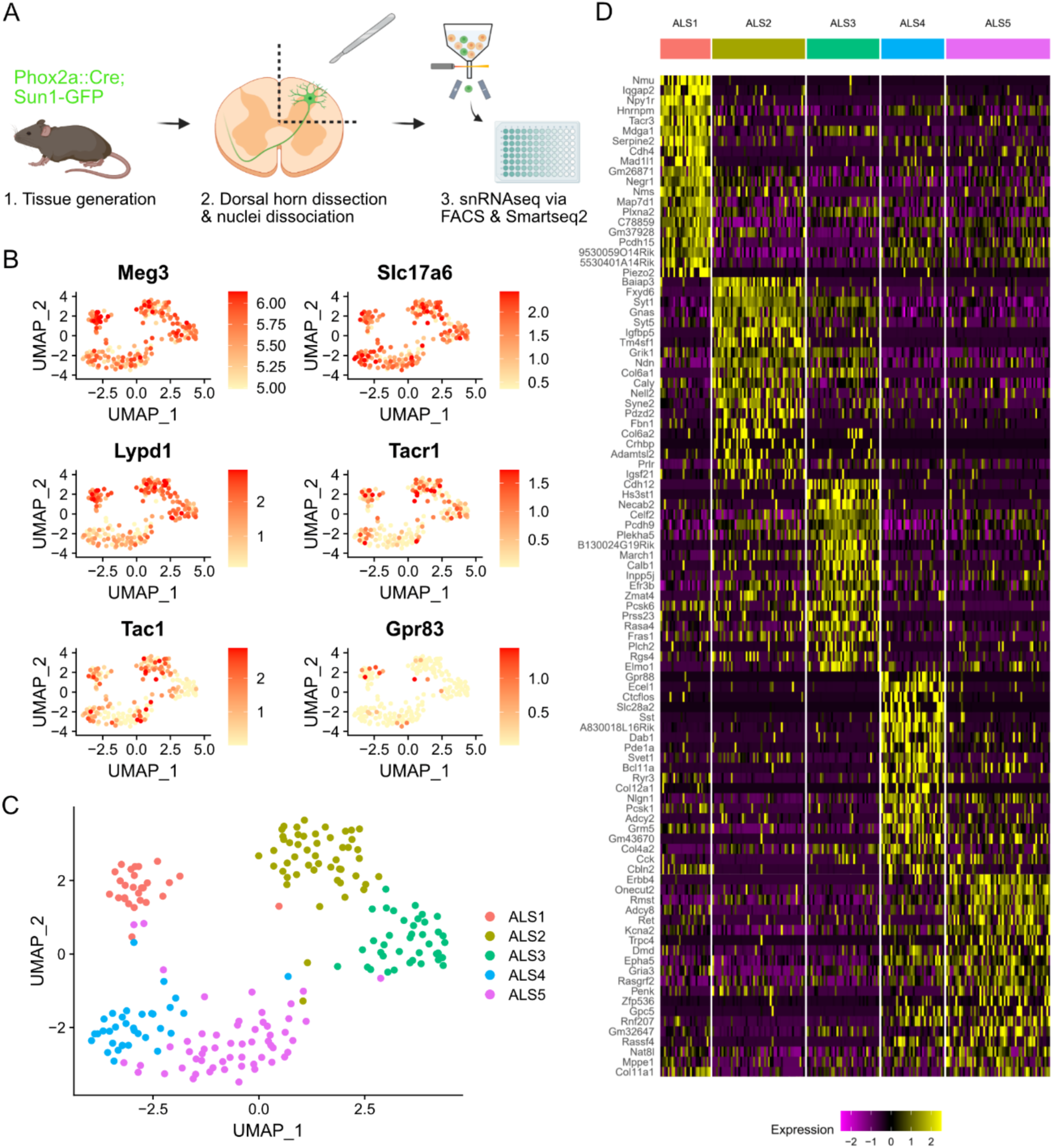
Transcriptomic clusters identified in the Phox2a::Cre mouse. (**A**) Schematic of the experimental workflow. (**B**) UMAP feature plots showing gene expression across 206 single nuclei included in the final analysis. *Meg3* and *Slc17a6* are markers of neurons and excitatory neurons, respectively. *Lypd1*, *Tacr1*, *Tac1* and *Gpr83* are previously proposed markers of ALS neurons or ALS neuron subsets. (**C**) UMAP plot showing the 5 clusters (ALS1-5) detected within Phox2a nuclei. (**D**) Heatmap showing the single-nucleus expression profile of 206 nuclei organised by the clusters ALS1-5. Twenty genes are shown per cluster, representing the top differentially expressed genes according to adjusted p-value.

Single Phox2a-GFP nuclei from the 16 animals were sorted into 96-well plates, with one nucleus per well, using a balanced plate layout that included nuclei from both sexes and conditions (SNI and Naïve) on each plate (Fig. S1D). Four plates were submitted for single-nucleus RNA sequencing and libraries were prepared using Smart-seq2 (Fig. S1E). Libraries were sequenced to a depth of approximately 1 million reads per nucleus. Reads were pseudoaligned using Kallisto (25) and further analysis was conducted using Seurat (26) in R. Initial quality control involved discarding data from nuclei with fewer than 5000 unique genes detected, and the removal of a small cluster of oligodendrocyte nuclei, based on expression of the *Mog* gene (Fig. S1F). Data from a total of 206 Phox2a-GFP nuclei, with a mean number of unique genes detected of 10,500 per nucleus, were included in further analysis (Fig. S1G). No major differences in quality metrics were evident between plates, sexes or conditions (Fig. S1H,I,J). Phox2a-GFP nuclei expressed high levels of the neuronal marker *Meg3* and the vesicular glutamate transporter *Slc17a6* (Fig. 1B). There was little to no expression of the inhibitory neuronal markers *Slc32a1* and *Gad1* (Fig. S1K). Therefore, the Phox2a-GFP nuclei sequenced here are all likely to be from excitatory neurons, as expected for cells belonging to the ALS. Many of the nuclei also expressed previously proposed markers of ALS neurons and ALS neuron subclasses such as *Lypd1*, *Tacr1* and *Tac1* (Fig. 1B & S2).

### ALS neuron subclasses are defined by unique molecular features

We first examined the gross structure of clusters based on UMAP projections and compared naïve and SNI, and male and female animals. We observed no difference in cluster structure between these groups (Fig. S3A,B). We also conducted a pseudo-bulk differential gene expression analysis to detect genes that may have been upregulated or downregulated in neuropathic pain specifically in nuclei of ALS cells. We did not detect any significant changes in gene expression following SNI (Fig. S3C,D). Based on these findings, all 206 nuclei were pooled for subsequent analysis.

Within the sequenced nuclei, 5 clusters were evident, and we named these ALS1-5 (Fig. 1C). Each cluster was defined by distinct patterns of gene expression (Fig. 1D). The ALS1 cluster is marked by expression of Neuromedin U (*Nmu*) but also, interestingly, showed selective expression of several other genes previously implicated in pain and sensory processing. These genes included neuropeptide Y receptor 1 (*Npy1r*), tachykinin receptor 3 (*Tacr3*), the α2a adrenoceptor (*Adra2a*) and neuropeptide FF receptor 2 (*Npffr2*). The ALS2 cluster was defined by expression of BAI1 Associated Protein 3 (*Baiap3*) and the *Fxyd6* gene. The expression of *Cdh12* and *Hs3st1* marked the ALS3 cluster, although *Cdh12* expression was also common, albeit at a lower level, in the ALS2 cluster. The ALS3 cluster also contained cells that express the calcium binding proteins *Necab2* and *Calb1*. Moreover, we noted a differential distribution of *Gria1* and *Gria4*, which code for the ionotropic glutamate receptor subunits GluA1 and GluA4, among the ALS1-3 clusters (Fig. S4A). The ALS4 and ALS5 clusters were closely related and shared the expression of several genes including the glycine receptor α1 subunit (*Glra1*) and the GABA_A_ receptor α1 subunit (*Gabra1*) (Fig. S4B). However, many genes were also differentially expressed in these clusters. The ALS4 cluster was marked by expression of *Gpr88* and *Ecel1* but also showed expression of the *Slc28a2* transporter, the peptide somatostatin (*Sst*), and corticotrophin-releasing hormone (*Crh*) (Fig. 1D & S2). Finally, the ALS5 cluster was defined by expression of *Erbb4* and *Onecut2*, and nuclei in this group also showed enriched expression of genes such as ionotropic glutamate receptor subunit 3 (*Gria3*), and proenkephalin (*Penk*).

The deep-sequencing approach employed here demonstrates the differential expression of transcripts across the separate clusters of Phox2a-lineage ALS neurons, and this may give insights into function. To better understand patterns of gene expression across clusters, we cross-referenced genes from the top 2000 variable features of our dataset with known databases of pain genes, peptides, G-protein linked receptors and both voltage-and ligand-gated ion channels. Dot plots showing the proportion of cells expressing each gene within each cluster and the normalised expression level were generated (Fig. S5). We also sought to align our single-nucleus data with existing sequencing data from the mouse dorsal horn. We integrated our data with cells sequenced by Haring et al. (16) and with a harmonised atlas of dorsal horn neurons (18). However, in both cases, our nuclei did not align closely to any class of cells (Fig. S6). There was no close relationship between our nuclei and the Glut15 (16) and Excit-21 (18) classes, respectively, which have been proposed as those that contain projection neurons. This was not unexpected due to the predicted rarity of ALS neurons within existing datasets, and to stark differences in technical approaches employed between these previous studies and ours.

### Spatial validation of ALS neuron clusters

The cell bodies of ALS neurons are present in multiple areas of the dorsal horn. They are concentrated in lamina I but are also scattered throughout deeper laminae. It has been suggested that the laminar location, in particular the distinction between deep and superficial cells, underlies differing functional roles of ALS neurons in pain (2, 27). We sought to confirm patterns of gene expression within the 5 clusters, and determined their laminar distribution, using multiplexed RNAscope *in situ* hybridisation (Fig. 2A). We selected a panel of 5 marker genes to distinguish the clusters: *Nmu*, *Baiap3*, *Cdh12*, *Gpr88* and *Erbb4*. These were chosen because their expression patterns were largely non-overlapping (Fig. S7A). Although these genes distinguished ALS clusters based on our sequencing findings, they are also widely expressed in other cells in the dorsal horn. The expression patterns of each marker across the dorsal horn are shown in Fig. S7B-F.

**Figure 2.**
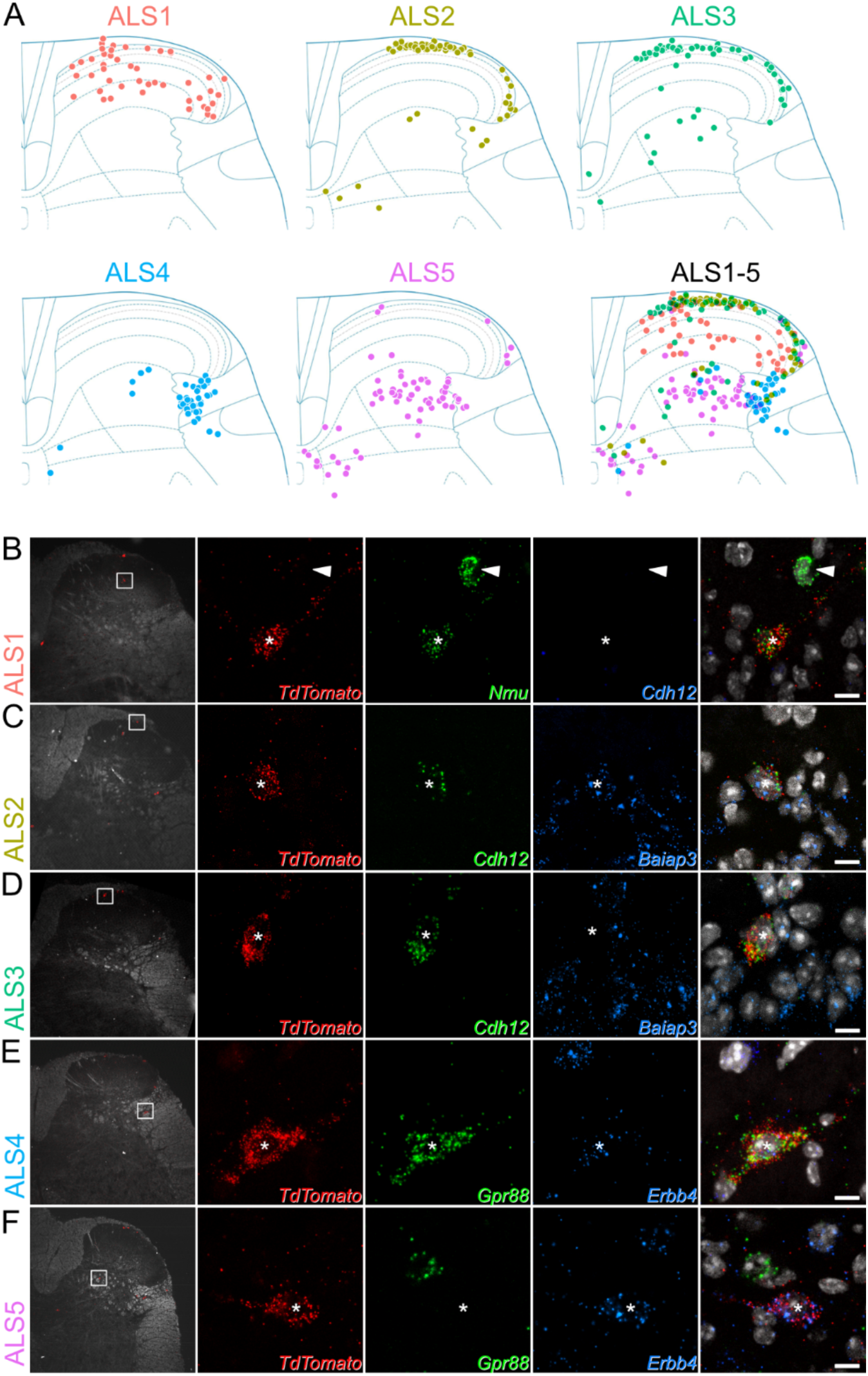
Laminar location of cells belonging to each cluster. (**A**) Plots showing the location of cells in each cluster. Each dot represents a single Phox2a cell with the appropriate *in situ* marker gene expression. Additional data to support cluster assignment is in Fig. S8. (**B**-**F**) *In situ* hybridisation signals seen with probes directed against *TdTomato* and mRNAs that were used to define each of the clusters in tissue from Phox2a::Cre;Ai9 mice. In each case the left pane shows a low-magnification view to indicate the location of the cell, with a box indicating the area seen at high-magnification in the remaining panes. The right pane is a merged image with each type of *in situ* hybridisation signal superimposed on nuclear staining (grey). Each set of high-magnification images contains a single *TdTomato*-positive Phox2a cell, which is marked with an asterisk. These cells are positive for *Nmu* (**B**), *Cdh12* and *Baiap3* (**C**), *Cdh12*, but not *Baiap3* (**D**), *Gpr88* and *Erbb4* (**E**) and *Erbb4*, but not *Gpr88* (**F**). The arrowhead in **B** shows another neuron that is positive for *Nmu* but not for *TdTomato*. High magnification scans are single confocal optical sections. Scale bar = 10 μm.

We conducted our analyses using lumbar spinal cord tissue from 4 Phox2a::Cre;Ai9 mice, in which Phox2a-lineage neurons are labelled with TdTomato (19, 21). We hybridised sections with probes against TdTomato and one of 3 separate marker pairs: *Nmu/Cdh12*, *Baiap3/Cdh12* and *Gpr88/Erbb4*. We used 12 dorsal horn sections per animal for each pair of markers. For each section, Phox2a cells were identified using low magnification confocal scans of the TdTomato channel (Fig. 2B-F). The laminar locations of the Phox2a cells were plotted onto outlines of the dorsal horn based on darkfield confocal scans that were acquired simultaneously. The transcriptional identity of each Phox2a cell was then recorded, based on higher magnification scans of the marker gene and TdTomato *in situ* hybridisation signals for each individual Phox2a neuron.

Phox2a neurons belonging to the ALS1 class were identified by the expression of *Nmu* (Fig. 2B). As predicted by the sequencing data, cells in this class did not co-express *Cdh12*. They represented 16% of all Phox2a neurons (Fig. S8A) and were distributed in lamina I, but also across laminae II-IV of the dorsal horn (Fig. 2A). The ALS2 and ALS3 clusters were identified using probes against *Baiap3* and *Cdh12* (Fig. 2C,D & S9). Cluster allocation was based on the highly specific expression of Baiap3 in ALS2 neurons, together with high expression of *Cdh12* in ALS3 and lower expression in some ALS2 cells (Fig. S7A & S8B). Based on our *in situ* analysis, cells in ALS2 and ALS3 each comprised 21% of all Phox2a neurons (Fig. S8B) and were located mainly in lamina I (Fig. 2A,C,D). The ALS4 and ALS5 clusters were identified *in situ* using probes against *Gpr88* and *Erbb4*, respectively (Fig. 2E,F). *Gpr88* was restricted to cells in ALS4, while *Erbb4* was found in all ALS5 cells and at lower levels in some of those belonging to ALS4. These clusters accounted for 16% (ALS4) and 26% of all Phox2a neurons counted (Fig. S8C). Cells belonging to both these clusters occupied the deeper laminae of the dorsal horn. ALS4 cells were highly concentrated in the lateral reticulated part of lamina V, with ALS5 cells scattered more broadly across the deeper laminae of the grey matter, both in lamina V and around the central canal (Fig. 2E,F). In summary, the identified neuronal types displayed distinct laminar distributions (Fig. 2A), suggesting that the location of ALS neurons is dependent on their transcriptional identity.

### The ALS1 cluster includes a previously recognised class of ALS neuron known as antenna cells

Based on the laminar distribution of cells within the ALS1 cluster, we suspected that these cells might include a previously identified class of ALS neurons known as antenna cells (28, 29), which are captured in the Phox2a::Cre mouse line (20). These cells are located in laminae II-V and have extensive dendritic trees that reach lamina I. Antenna cells are also distinctive as they receive dense synaptic input from bundles of peptidergic nociceptors (20, 30).

We tested the hypothesis that cells belonging to the ALS1 cluster included antenna cells by using a combined *in situ* hybridisation and immunohistochemistry approach (ISH/IHC). In sagittal sections of lumbar spinal cord from 3 Phox2a::Cre;Ai9 mice, we identified antenna cells by immunostaining for TdTomato and the peptidergic nociceptor marker CGRP. This was conducted alongside *in situ* hybridisation with a probe against *Nmu* (Fig. 3). As expected, *Nmu* was detected in a large number of lamina II interneurons (16). In this tissue we identified 29 (26–34) antenna-type ALS cells per animal, based on the characteristic pattern of contacts from CGRP-immunoreactive axons (20). The vast majority of these (94%, 92-97%) also expressed *Nmu*. In addition, we identified a small number of non-antenna ALS cells in lamina I that expressed *Nmu*. We counted 15 (11–19) of these cells per animal, representing only 11% (9-13%) of all lamina I Phox2a-TdTomato cells. Antenna cells outnumbered the lamina I *Nmu*+ Phox2a-Tdtomato cells by a ratio of 1.9:1 (1.7-2.2-fold). We also observed qualitatively that the *Nmu*+ lamina I ALS cells were located deeper in this lamina than other Phox2a-Tdtomato cells, and that they often had ventrally directed dendrites. In some cases, these were associated with CGRP axons (12) (Fig. S10). These data suggest that the ALS1 cluster includes the previously identified class of ALS neurons known as antenna cells, and that these cells have a rarer transcriptionally-related counterpart in lamina I.

**Figure 3.**
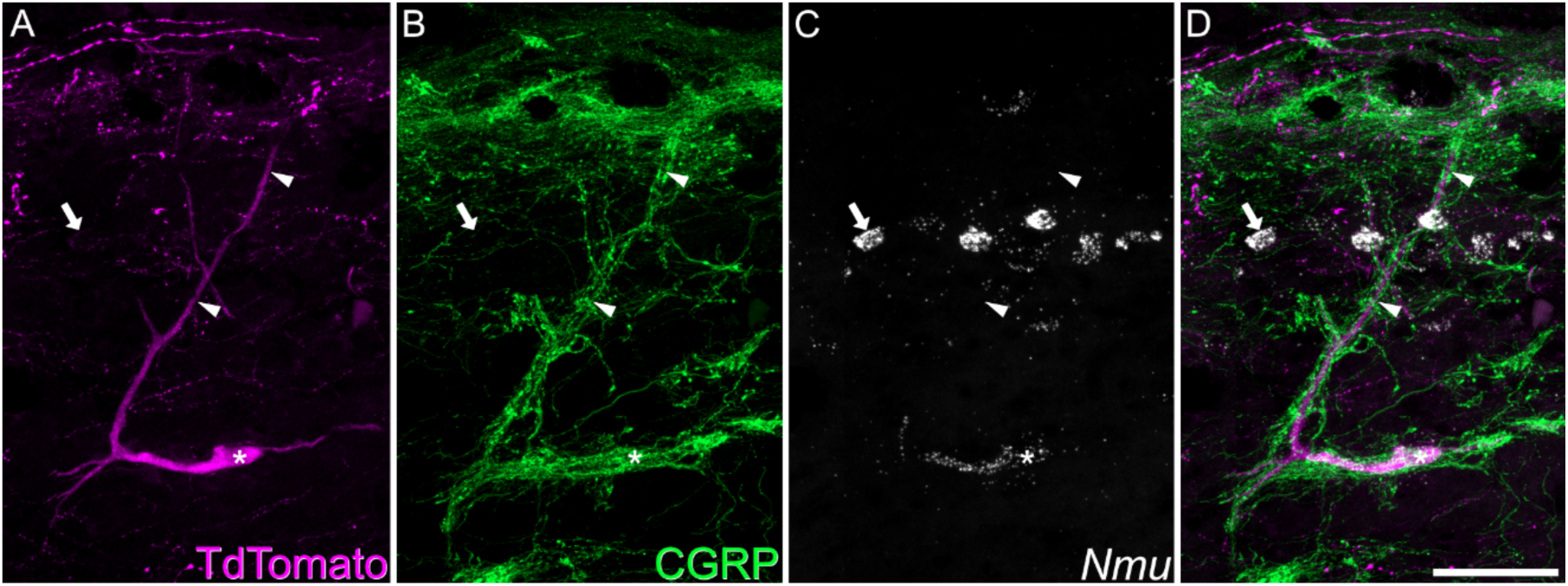
Expression of *Nmu* by a Phox2a+ antenna cell. Combined *in situ* hybridisation and immunohistochemistry performed on a sagittal section of lumbar spinal cord from a Phox2a::Cre;Ai9 mouse. **A** and **B** show immunostaining for TdTomato and CGRP, while **C** shows *in situ* hybridisation signal for *Nmu*, and **D** is a merged image. A large TdTomato-labelled cell is indicated with an asterisk. It has a long dorsal dendrite (marked with arrowheads) that is surrounded by numerous CGRP-immunoreactive axons, indicating that this is an antenna cell. *In situ* hybridisation shows that this cell expresses *Nmu* mRNA, which is also present in many small interneurons in lamina II (one shown with an arrow). This image is a projection of confocal optical sections (1 μm z-separation) taken through the full 12 μm thickness of the section. Scale bar = 50 μm.

### Cells within the ALS3 cluster receive preferential input from TRPM8 expressing thermoreceptors and include putative ‘cold-selective cells’

There is considerable evidence that a subpopulation of lamina I ALS cells respond selectively to skin cooling (9, 11–14), and using a semi-intact preparation (31), we have found that around a third of Phox2a-positive lamina I projection neurons respond only to cold stimuli applied to the skin (JH, unpublished observations). We looked for an anatomical approach to identify these cells that could be combined with the detection of molecular markers through *in situ* hybridisation. The cold-selective cells are thought to be innervated by cold-sensing primary afferents that express the TRPM8 channel (11). We therefore injected AAV.mCherry into the LPb of Trpm8^Flp^;RCE:FRT mice, in which TRMP8 afferents express GFP. In horizontal sections we noted that TRPM8 afferents formed interweaving bundles in lamina I, and that these did not occupy the entire area of the lamina. We found that some mCherry-labelled lamina I ALS neurons received numerous contacts from TRPM8 afferents. For these cells, the afferents typically surrounded the cell body and extended along the dendrites (Fig. 4A). In the L4 segments of 4 mice the neurons with dense innervation from TRPM8 afferents accounted for 21% (17.6-29.7%) of all mCherry-labelled lamina I ALS cells. We also observed that the remaining retrogradely labelled lamina I neurons received relatively few or no contacts from TRPM8 boutons. To confirm the dichotomy between the cells with dense TRPM8 input and the remaining lamina I projection neurons, we quantified the proportion of excitatory synaptic input from TRPM8 afferents onto ALS cells. We reconstructed a total of 20 lamina I ALS neurons, 10 of which had been identified as densely innervated, and 10 as not densely innervated. We identified an average of 159 excitatory synapses (range 43-320) per cell, by examining immunostaining for the post-synaptic density protein Homer (Fig. S11). For those cells classified as densely innervated, the proportion of excitatory synapses originating from TRPM8 (GFP-positive) boutons was 62%, and this was considerably higher than for cells without dense TRPM8 innervation, for which this figure was only 6% (Fig. S11A). Furthermore, by generating the more complex Trpm8^Flp^;RCE:FRT;Phox2a::Cre;Ai9 cross, we were able to confirm that Phox2a-positive lamina I cells were included among those that were densely innervated by TRPM8 afferents (Fig. S12).

**Figure 4.**
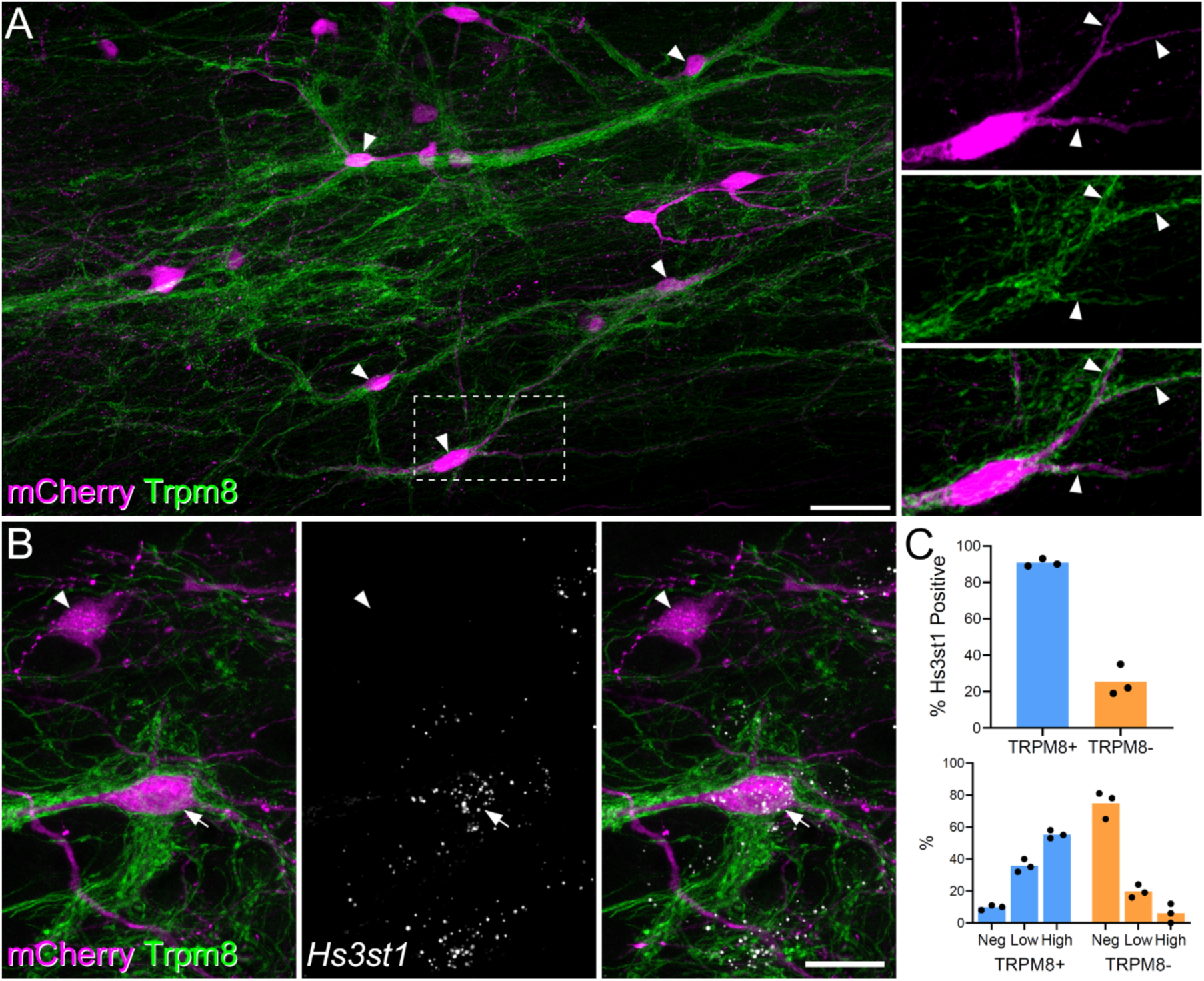
Projection cells with dense Trpm8 input are likely to correspond to ALS3. (**A**) Horizontal section through lamina I showing dense innervation of spinoparabrachial neurons (retrogradely labelled with AAV.mCherry injected into the lateral parabrachial area, LPb) by GFP-labelled axons in a Trpm8^Flp^;RCE:FRT mouse. Arrowheads indicate mCherry-positive cells that receive numerous contacts from GFP-labelled (TRPM8) axons. The box indicates the area that is enlarged to the right of the main image. This shows staining for mCherry and GFP separately and in a merged view. Note that the cell body and dendrites (3 indicated with arrows) are associated with GFP-labelled axons. Main and inset images are maximum projection of 63 and 16 (respectively) optical sections at 0.5 μm z-spacing. Scale bar = 50 μm. (**B**) Combined immunohistochemistry and fluorescent *in situ* hybridisation on a horizontal section through lamina I from another Trpm8^Flp^;RCE:FRT mouse that had received an injection of AAV.mCherry into the LPb. Two retrogradely labelled (mCherry-positive) cells are seen. One of these (arrow) is densely coated with GFP-labelled (TRPM8) axons, while the other (arrowhead) is not. The cell with numerous TRPM8 contacts contains *Hs3st1* mRNA, whereas the other cell lacks this mRNA. This image is a projection of confocal optical sections (1 μm z-separation) taken through the full 12 μm thickness of the section. Scale bar = 20 μm. (**C**) Quantification of *Hs3st1* transcripts on mCherry cells with dense TRPM8 innervation (blue bars, TRPM8+) and mCherry cells that lacked dense TRPM8 innervation (orange bars, TRPM8-). Each dot represents data from a single mouse.

Hachisuka et al (11) reported that the cold-selective ALS cells responded weakly, if at all, to substance P, suggesting that they express low levels of mRNA for the cognate receptor, *Tacr1*. Among the two remaining lamina I clusters (ALS2 and ALS3), cells in ALS3 showed a lower incidence of *Tacr1* expression (Fig. 1B), and we therefore asked whether these neurons corresponded to those with dense TRPM8 input. To test this, we analysed tissue from 3 Trpm8^Flp^;RCE:FRT mice that had received injections of AAV.mCherry into the LPb. Horizontal sections from lamina I were reacted with antibodies against GFP and mCherry, following *in situ* hybridisation with probes against *Hs3st1*, as a marker of ALS3 neurons. For these experiments we chose *Hs3st1* (rather than *Cdh12*) to identify ALS3 cells as it is highly restricted to cells in this cluster.

We identified an average of 93 (range 64-133) mCherry+ lamina I ALS neurons per animal. Each cell was graded with respect to TRPM8 inputs by an assessor who was unaware of the *Hs3st1* transcript levels. Cells that had a sufficient length of dendrite in the section were graded as densely TRPM8 innervated or not densely innervated. *Hs3st1* transcript levels per cell were then recorded as absent, low (5-15 transcripts per cell) or high (>15 transcripts per cell) by a second observer, who was unaware of the classification by TRPM8 input. We identified 26 (19–40) neurons with dense TRPM8 input per animal, representing 27% (23-31%) of all mCherry+ ALS cells in lamina I, and found that 91% (89-93%) of these cells were positive for *Hs3st1* (Figs. 4B,C). Among those cells that lacked dense TRPM8 input only 25% were positive for *Hs3st1* and notably, the expression level of Hs3st1 in these cells was lower (Fig. 4C). Taken together, these findings strongly suggest that the ALS3 cluster includes the cold-selective lamina I projection neurons identified in previous studies.

### Molecular-genetic retrograde tracing allows targeting of specific clusters of ALS neurons

Finally, we asked whether subpopulations of ALS neurons could be targeted selectively by using mouse genetics and retrograde viral tracing (Fig. 5). Mouse lines in which Cre-recombinase expression is driven by the marker gene of interest received injections of AAVs encoding Cre-dependent constructs into the LPb to specifically label subsets of projection neurons. We used two well-established lines in which Cre is knocked into either the somatostatin or the cholecystokinin locus (Sst^Cre^ and Cck^Cre^) (32).

**Figure 5.**
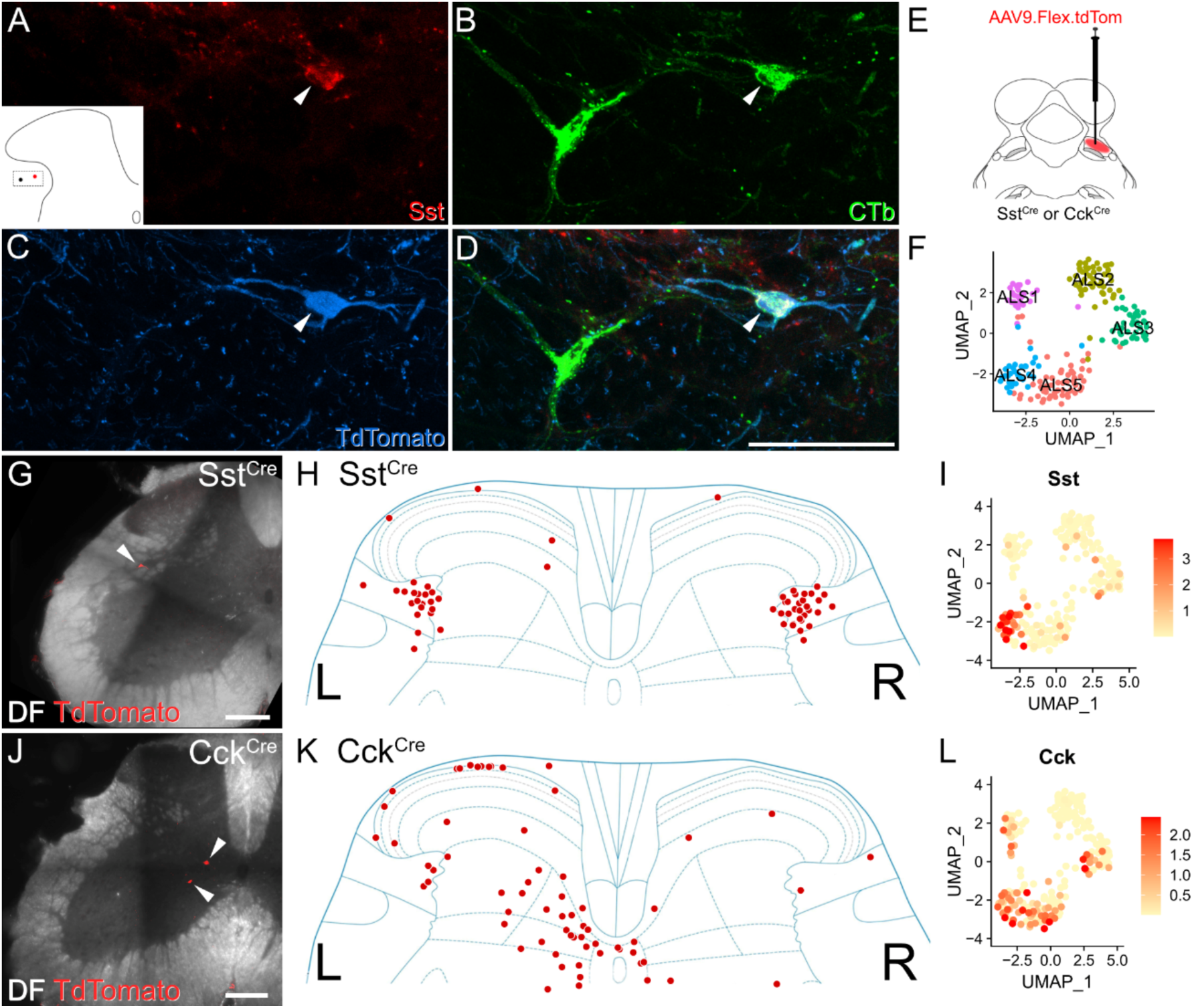
Retrograde viral tracing of ALS cells. (**A**-**D**) Immunohistochemistry showing the presence of somatostatin (Sst) in ALS neurons in lateral lamina V. The lateral lamina V area of a lumbar transverse spinal cord section from a Phox2a::Cre;Ai9 that had received an injection of CTb into the LPb is shown. The section has been stained for Sst, CTb and TdTomato, and these are shown separately in **A**-**C** and merged in **D**. Two CTb labelled ALS neurons are present, and the location of these is shown in the inset in **A**. One of these is a Phox2a cell (marked with an arrowhead) and is also positive for Sst. Images are a maximum projection of 6 optical sections at 1 µm z-spacing. Scale bar = 50 μm. (**E**-**L**) Retrograde labelling of molecularly defined subclasses of ALS neurons. (**E**) A schematic of the approach used. AAV.flex.TdTomato was injected into the right LPb of either Sst^Cre^ or Cck^Cre^ mice (n = 2 for each genotype). This resulted in labelling of Cre-expressing ALS cells in the lumbar spinal cord. (**F**) UMAP plot showing the 5 clusters. Representative images of retrogradely labelled cells are shown in **G** and **J**. In Sst^Cre^ animals, traced cells were seen mainly in reticulated lateral lamina V bilaterally as shown in **H**. Cells traced in Cck^Cre^ animals were present across the superficial and deep dorsal horn and are mainly contralateral to the injection site (**K**). In both cases, these distributions are consistent with the pattern of expression of the mRNAs across the clusters, as shown in UMAP plots **I** and **L**.

Although somatostatin is widely expressed in excitatory interneurons in the dorsal horn of the spinal cord (33, 34), our data shows that it is also seen at high levels in the ALS4 cluster (Figs. 1D & 5F,I). These cells have a restricted distribution, with cell bodies in lateral lamina V (Fig. 2A). In order to verify that somatostatin was expressed in ALS cells in this region, we first performed immunohistochemistry for the peptide in spinal cord sections from 2 Phox2a::Cre;Ai9 mice. These animals had received injections of cholera toxin subunit B (CTb) into the LPb, thereby allowing identification of both Phox2a-positive and Phox2a-negative projection neurons (19). We observed somatostatin labelling in 54% of CTb-labelled lateral lamina V projection neurons (Fig. 5A-D), and a similar proportion of those within the Phox2a subset (48%) (Table S1). This proportion was considerably higher than for projection neurons in other areas of the cord in which somatostatin immunoreactivity was observed in approximately 9% of cells (Table S1).

We then injected AAV.flex.TdTomato into the right LPb of two Sst^Cre^ mice, to label somatostatin-expressing ALS neurons (Fig. 5E). As predicted, we found that the cell bodies of these neurons were located predominantly in lateral lamina V. Interestingly, similar numbers of retrogradely labelled neurons were found on the sides contralateral and ipsilateral to the brain injection (Fig. 5G,H). This suggests that the supraspinal projection pattern of somatostatin-expressing ALS neurons differs from that of most other ALS cells, which project mainly to the contralateral side (1, 2, 6).

We also compared the distribution of retrogradely labelled somatostatin-positive ALS neurons with those labelled using the same approach in a CCK^Cre^ mouse line. In contrast to *Sst*, *Cck* is found in Phox2a cells in most clusters, apart from ALS2 (Figs. 1D & 5F,L). As expected (based on the laminar distribution of cells in ALS1 and ALS3-5; Fig. 2A), TdTomato-labelled neurons were located throughout the dorsal horn and these cells were far more numerous on the side contralateral to the LPB injection (Fig. 5J,K). In summary, our findings based on molecular genetic retrograde tracing support our sequencing and *in situ* hybridisation data by recapitulating predicted cluster distributions *in vivo*.

## Discussion

Our main findings are that ALS projection neurons belonging to the Phox2a-lineage can be divided into 5 discrete transcriptomic clusters, and that these are differentially distributed across the dorsal horn. Two of these clusters, ALS1 and ALS3, appear to include well-recognised functional populations, antenna cells in laminae II-V and cold-selective lamina I neurons, respectively. We also provide a comprehensive molecular dataset that will allow identification of genes of interest across distinct ALS subclasses.

### Lack of evidence for changes following nerve injury

In first order neurons, peripheral nerve injury causes a dramatic transcriptional response, leading to loss of cell identity (35). There is also evidence for more modest changes affecting spinal projection neurons (36, 37). We therefore took the opportunity to look for altered gene expression in projection neurons in the L4 segment, which is within the region affected by SNI. However, we did not find any differences, either in gross cluster structure or in pseudo-bulk gene expression, between conditions. The study was not powered to look for transcriptional changes in each of the clusters individually, or indeed to detect small fold-changes in lowly expressed genes (e.g., GPCRs). However, now that we have identified clusters among the ALS cells, it may be possible in future to isolate distinct cell types and definitively address whether transcriptional changes occur in neuropathic pain states.

### Phox2a-lineage ALS cells

Developmentally, the transcription factor Phox2a is expressed by a subset of those dI5-lineage spinal neurons that project to the brain (17). Transient expression of Phox2a therefore defines a population of projection neurons, and importantly, cells captured by the Phox2a::Cre line are thought to belong exclusively to the ALS (19–21). However, several groups of ALS neurons are under-represented. Specifically, the Phox2a::Cre;Ai9 cross captures up to 60% of the those in lamina I, most (if not all) of the antenna cells, but very few of those in the LSN. In the deep dorsal horn, up to 25% of spinoparabrachial cells are labelled (19–21). The ALS cells that are not detected in crosses involving the Phox2a::Cre mouse may belong to other populations that do not depend on Phox2a expression, or result from incomplete capture of Phox2a-lineage neurons in this BAC transgenic line. It is therefore difficult to predict the extent to which the classification scheme proposed here would extend to the entire ALS. It is very likely that non-Phox2a lineage cells would belong to different clusters. However, it is also conceivable that despite their different developmental origins, ALS neurons undergo a convergence of transcriptional phenotype in the adult, such that these would map onto the clusters that we have identified.

Although cells belonging to 4 of the clusters (ALS1-4) show a relatively restricted spatial distribution, ALS5 includes cells in the lateral part of lamina V as well as those near the central canal, and these presumably represent functionally discrete populations. Sequencing a larger number of cells may therefore reveal further subdivisions, particularly among cells in ALS5. Overall, it is likely that future studies will reveal an even greater number of discrete subpopulations within the ALS, reflecting the functional diversity of this system.

### Relation to previously defined ALS neurochemical populations

The neurokinin 1 receptor (NK1r), encoded by *Tacr1*, has long been regarded as a marker for a large subset of ALS neurons, particularly those in lamina I (38, 39). However, its usefulness for defining functional subsets has been questioned, due to its widespread expression among projection neurons (2). Consistent with this, we show that *Tacr1* is expressed to varying degrees among each of our clusters, albeit at lower levels in ALS3 and ALS5. More recently, Choi et al (23) proposed that lamina I ALS cells could be assigned to largely non-overlapping classes defined by expression of *Tacr1* or *Gpr83*. Although we see *Gpr83* in each of our clusters, cells with this mRNA are infrequent, presumably due to low levels of expression, and the failure to detect these transcripts with nuclear sequencing. Another gene that has been identified in subsets of ALS neurons is *Tac1*, which was found in ALS neurons in several regions of the dorsal horn (22, 40). Consistent with this, we find *Tac1* expressed in all of our clusters.

As far as we are aware, the only previous study to employ RNA sequencing to investigate heterogeneity within the ALS was that of Wercberger et al (24), who used a retro-TRAP (translating ribosome affinity purification) technique to identify candidate projection neuron genes. They noted an extensive molecular diversity of spinoparabrachial neurons, and proposed *Cck*, *Nptx2*, *Nmb* and *Crh* as markers of distinct molecular subsets. We were able to demonstrate expression of all four of these mRNAs in our dataset, with certain clear patterns in relation to the clusters. Although expression of *Nptx2* or *Nmb* was seen broadly across our clusters, both *Cck* and *Crh* had a more restricted distribution. Specifically, *Cck* was present in all clusters apart from ALS3, and this is consistent with the *in situ* hybridisation data of Wercberger et al (24), which showed expression in both deep and superficial laminae. *Crh* displayed a more restricted pattern, being present mainly in the ALS4 cluster, with a few cells in ALS2.

### Unbiased clustering as a means to define functional populations within the ALS

Somatosensory information is processed through circuits involving primary afferents, dorsal horn interneurons and ALS projection neurons (6). Recent advances in the classification of the afferents (41, 42) and interneurons (16, 18) by unbiased RNA sequencing have revolutionised our understanding of these components. However, despite the known heterogeneity of ALS cells, until now we have lacked equivalent insight into the molecular architecture of this system. In turn, this has hindered attempts to define the complex synaptic circuitry underlying pain, itch and temperature sensation.

The functional relevance of the classification scheme proposed here is supported by the spatial separation between cells belonging to the clusters, as well as by the alignment of two of the clusters (ALS1 and ALS3) with previously identified functional populations. Most cells in ALS1 were located in laminae II-IV, and based on their location and their close association with peptidergic nociceptors, these were identified as antenna cells. These cells are remarkably homogeneous in both their morphology and synaptic inputs. In addition to receiving input from peptidergic nociceptors, they are also densely innervated by NPY-expressing inhibitory interneurons (20, 30, 43, 44). Interestingly, cells in this cluster showed robust expression of the NPY1 receptor (*Npy1r*), suggesting a microcircuit with a possible role in NPY-mediated analgesia (45). It has been proposed that both antenna cells and lamina I Phox2a cells share a common birthdate and migration pattern (21). Our finding that antenna cells have a transcriptomic counterpart in lamina I raises the possibility that the ALS1 cells in lamina I originate from the same population, but have failed to migrate ventrally.

The cold-selective cells, which are likely to be included in our ALS3 cluster, are a well-recognised population among lamina I ALS neurons (9, 11, 13, 14), and may represent a labelled line for cold sensation (46). With the exception of these cells, ALS neurons in lamina I are thought to respond to a variety of thermal and mechanical noxious stimuli (2, 9), and the ALS2 population is therefore likely to consist mainly of nociceptive-specific cells.

We previously identified differential expression patterns for synaptic AMPA receptor subunits on ALS neurons in laminae I-IV, with GluA1 and GluA4 subunits showing a non-overlapping distribution at excitatory synapses on these cells (47, 48). GluA4 (encoded by *Gria4*) was expressed by the antenna cells and by large lamina I ALS cells, whereas GluA1 (encoded by *Gria1*) was found exclusively on smaller ALS cells in lamina I. Consistent with this, we find that cells in ALS1 (which includes the antenna cells) showed relatively strong expression of *Gria4*. This also suggests that the lamina I component of ALS1 includes some of the large lamina I ALS cells that express GluA4. Cells in ALS2 and ALS3 show relatively high *Gria1* expression and probably correspond to the smaller lamina I ALS cells with synaptic GluA1 receptors.

Compared to superficial ALS neurons, much less is known about those in deeper laminae. In contrast to the predominantly nociceptive-specific cells in lamina I, these cells are thought to have wide dynamic range (WDR) receptive fields, responding to both noxious and innocuous stimuli (27), and activity in these deep ALS neurons is sufficient to evoke pain in humans (49, 50). Notably, our analysis clearly separates these from lamina I and antenna ALS cells, and reveals two separate classes, ALS4 and ALS5. Certain genes were differentially expressed between these and the more superficial clusters. *Erbb4* was present in both of the deep clusters, consistent with the findings of Wang et al (51), who identified Cre+ projection cells in the deep (but not superficial) dorsal horn in Erbb4^CreERT2^ mice. Glycine, rather than GABA, is the main fast inhibitory transmitter in deep dorsal horn (52), and we found that *Glra1* (which codes for the glycine receptor α1 subunit) was largely restricted to cells in ALS4 and ALS5. The GABA_A_ receptor α1 subunit is excluded from the SDH (53), and the corresponding mRNA (*Gabra1*) was seldom seen in the ALS1-3 clusters.

Most cells in the ALS4 cluster were found in a lateral region corresponding to the reticulated part of lamina V (54). Many of these cells expressed *Sst* and this is of interest given the proposed role of somatostatin-expressing spinal neurons in mechanical pain and pruritogen-evoked itch. Duan et al. reported that ablation of somatostatin-expressing spinal neurons resulted in loss of acute mechanical pain, and a dramatic reduction of both dynamic and static mechanical allodynia in neuropathic and inflammatory models (33). In a subsequent study it was reported that this ablation significantly attenuated scratching evoked by a variety of pruritogens (55). These findings were attributed to loss of excitatory interneurons in SDH, most of which express somatostatin (34). However, our finding that *Sst* is also expressed in the ALS4 cluster raises the possibility that loss of projection neurons may have contributed to the behavioural phenotype seen following ablation of somatostatin-expressing cells.

Overall, our results provide important novel insights into the molecular logic of the ALS, which will facilitate targeting of specific populations and manipulation of their activity *in vivo*. Combining these approaches with recent advances in the study of mouse behaviour (56–58), should allow resolution of longstanding controversies, such as the existence of labelled lines, and the relative contribution of deep versus superficial projection neurons to different aspects of the pain experience (1–3).

## Materials and Methods

All experiments were carried out in accordance with the UK Animals (Scientific Procedures) Act 1986 and adhered to ARRIVE guidelines. Transgenic mice, experimental procedures, viruses, antibodies and probes and are described in SI Appendix, Materials and Methods.

## Data, Materials and Software Availability

Sequencing data are available to browse via the Broad Institute single cell portal at https://singlecell.broadinstitute.org/single_cell/study/SCP2337. Raw sequencing data, aligned counts, and the Seurat object are available to download from the Gene Expression Omnibus under the accession number GSE240528. This study did not generate new unique reagents or software.

## Acknowledgments

This research was funded in whole, or in part, by the Wellcome Trust (Grant numbers 219433/Z/19/Z and 204820/Z/16/Z), the Medical Research Council (Grant numbers MR/T01072X/1, MR/V033638/1, MR/W004739/1 and MR/W002426/1), the Biotechnology and Biological Sciences Research Council (BB/S017178/1), the Academy of Medical Sciences (Grant number SGL025\1079) and the Medical Research Foundation (MRF-160-0015-ELP-DENK-C0844). The authors acknowledge support from the Biotechnology and Biological Sciences Research Council (BBSRC), part of UK Research and Innovation, Core Capability Grant BB/CCG1720/1and the National Capability Grant BBS/E/T/000PR9816. This work was supported by the intramural research program of the NIDCR, NIH, project ZIADE000721-22. We acknowledge the assistance of Diane Vaughan, School of Infection and Immunity Flow Cytometry Facility at the University of Glasgow. We are grateful to Brian Roome and Sara Villa-Hernandez for helpful discussion, to Oscar Marin Parra and Eleanor Paul for sharing Cck^Cre^ and Sst^Cre^ mice, to Ariel Levine for sharing data, and to Robert Kerr and Iain Plenderleith for expert technical assistance. For the purpose of Open Access, the authors have applied a CC BY public copyright licence to any Author Accepted Manuscript version arising from this submission.

