## Supplemental Information for "Deep sequencing of Phox2a nuclei reveals five classes of anterolateral system neurons"

**This PDF file includes:**

Supporting methods
Figures S1 to S12
Table S1-3
SI References

### Supporting Methods

#### Animals & operative procedures

All experiments were carried out in accordance with the UK Animals (Scientific Procedures) Act 1986 and adhered to ARRIVE guidelines.

To identify Phox2a-lineage ALS neurons we used the BAC transgenic Phox2a::Cre mouse line in which Cre recombinase is expressed under control of the Phox2a promoter. This was crossed with different reporter lines: (CAG-Sun1/sfGFP or Ai9(RCL-tdT)), in which Cre-mediated excision of a STOP cassette drives expression of either superfolder GFP conjugated to the Sun1 nuclear membrane protein (CAG-Sun1/sfGFP), or TdTomato (Ai9). Crosses with the Sun1-GFP mouse line were used to extract nuclei for sequencing, whereas the Ai9 cross was used for identification of Phox2a-lineage cells using *in situ* hybridisation and immunohistochemistry. In the Phox2a::Cre;Sun1-GFP animals that underwent spared nerve injury (SNI), we ligated and transected the tibial and common peroneal branches of the sciatic nerve, leaving the sural nerve intact (59). Animals received postoperative analgesia comprising buprenorphine (0.5 mg/kg) and carprofen (5 mg/kg). These animals were euthanised and spinal cord tissue for sequencing was collected 8 days after the SNI procedure.

We also used Trpm8$^{Flp}$ mice crossed with the RCE:FRT reporter in which Flp mediated excision of a STOP cassette results in EGFP expression in Flp containing cells. Sst-IRES-Cre and Cck-IRES-Cre mice were used in retrograde tracing experiments. Injections into the LPb were performed as described previously (19). Archived tissue from the two Phox2a::Cre;Ai9 animals that had received bilateral CTb injections into the LPb in a previous study (19) was used here for immunohistochemical quantification of the proportion of ALS neurons expressing somatostatin. Trpm8$^{Flp}$;RCE:FRT mice, Sst$^{Cre}$ mice and Cck$^{Cre}$ mice received injections of viruses into the LPb as part of this study. The Trpm8$^{Flp}$ mice received right-sided unilateral LPb injections of AAV9.CB7.CI.mCherry.WPRE.rBG and were perfused with fixative 14 days following the injection. Sst$^{Cre}$ and Cck$^{Cre}$ mice received unilateral injections of ssAAV-1/2-CAG-dlox-tdTomato(rev)-dlox-WPRE-bGHp(A) into the LPb and were perfused 16-20 days after injection. In all cases of viral injection, the volume used was 1000 nl.

All mice were adults aged 9 to 16 weeks old, with the exception of Sst$^{Cre}$ and Cck$^{Cre}$ mice (18-27 weeks old), and were housed on a 12 h light/dark schedule.

#### Nuclei isolation and fluorescence-activated cell sorting

Phox2a::Cre;Sun1-GFP mice (n = 16) were deeply anaesthetised with isoflurane and decapitated. The lumbar spinal cord was rapidly removed and the dorsal half of the left side of the L4 segment dissected and frozen on dry ice. Tissue was stored at -80 °C for up to 1 month before further processing.

In preliminary experiments, we found that cell lysis using a mechanical-detergent technique significantly reduced the yield of Phox2a nuclei and therefore nuclei were extracted from the frozen spinal cord tissue using a modified hypotonic lysis protocol (60). Tissue from each mouse was processed separately as follows. The frozen spinal cord tissue was defrosted, chopped into small pieces and placed into 5 ml of hypotonic lysis buffer in a 15 ml centrifuge tube on ice. The hypotonic lysis buffer contained Tris-HCl (pH 7.4) 10 mM, NaCl 10 mM, MgCl 3 mM, and Nonidet P40 0.01% in nuclease free water. Tissue was left to stand on ice for 15 minutes and then 5ml of Neurobasal medium (2% B27 supplement, 1% N2 supplement, 1% Glutamax, (Thermo Fisher Scientific)) was added. The tissue was triturated using a series of 4 progressively smaller diameter fire polished pipettes. Each trituration involved passing the tissue in and out of the pipette ten times until it flowed easily. After each round, 2ml of supernatant was removed and passed through a 70 µm strainer into another 15ml centrifuge tube. Following the fourth round of trituration, after which tissue was no longer visible, the remaining liquid was passed through the filter into the second tube. The solution of dissociated nuclei was then centrifuged at 1,000 x g for 10 minutes at 4 °C. The resulting pellet was resuspended in 2.5 ml of low sucrose buffer (Sucrose 0.32 M, HEPES (pH = 8.0) 10 mM, CaCl 5 mM, MgAc 3 mM, DTT 1mM, EDTA-free Protease Inhibitor Cocktail (Roche) 1 tablet per 10 ml, RNAsin RNAase inhibitor (Promega) 40 U/ml, all in nuclease free water). A 2.5 ml sucrose cushion (Sucrose 1 M, HEPES (pH = 8.0) 10 mM, MgAc 3 mM, DTT 1 mM, all in nuclease free water) was then layered underneath using a serological pipette and the tube centrifuged at 3,200 x g for 20 minutes at 4 °C. Following centrifugation, the supernatant was discarded, and the pellet resuspended in 0.5 ml of 1x PBS containing 2% BSA, EDTA-free Protease Inhibitor Cocktail (Roche) (1 tablet per 10 ml) and RNAsin 40 U/ml. The solution was transferred to a 6 ml FACS tube and kept on ice prior to FACS.

FACS was conducted on a BD FACS Aria IIU equipped with a 100-micron nozzle, and BD FACSDiva software version 9.01. Prior to sorting, nuclei were stained with DRAQ7 (1:500, Invitrogen) to enable the sorting of nuclei from debris. Phox2a-lineage nuclei were identified based on GFP fluorescence and sorted into 96-well LoBind plates (Eppendorf) using single-cell purity settings such that each well contained a single nucleus. Prior to sorting, each well of the plate was filled with 2 µl of nuclease free water containing Triton X-100 0.2% and RNAsin 1 U/µl. The plate layout is shown in Fig S1, and all four plates submitted for sequencing were filled over two FACS sessions on two separate days. Plates 1 and 2 on day 1 and plates 3 and 4 on day 2. Once filled with nuclei, plates were sealed, spun briefly (1,000 x g for 1 minute) and flash frozen at -80 °C. Control samples were also run as part of each FACS session to verify the gating strategy. Spinal tissue from Sun1-GFP mice without any Cre-expressing alleles (i.e. non-Phox2a::Cre) was processed as above. This tissue was split into two FACS tubes, with one stained with DRAQ7 and one remaining unstained. Very few events were observed in the nuclei gate in the absence of DRAQ staining, and no events were seen in the GFP gate in the absence of Cre-expression.

**Single cell RNA sequencing-library construction**

Plates were stored at -80 °C until further processing according to standard Smart-seq2 protocol (61). Reverse transcription conditions were: 42°C for 90 minutes, 10 cycles 50°C for 2 min, 42°C for 2 min, hold at 4°C; PCR amplification conditions were: 98°C for 3 min, 21 cycles: 98°C for 20 sec, 67°C for 15 sec, 72°C for 6 min, final extension 72°C for 5 min, hold at 4°C. Amplified cDNA underwent bead clean-up in a 1:0.8X volumetric ratio of AMPure XP Beads (Beckman Coulter) and eluted into 25 µl of Nuclease Free Water. Selected libraries were analysed on the Bioanalyzer using a High Sensitivity DNA chip (Perkin Elmer) to assess their quality. Sequencing libraries were generated using an automated, reduced-volume version of the Nextera XT protocol (Illumina) using the SPT Labtech Mosquito LV. Libraries were cleaned up using a 1:0.7x volumetric ratio of AMPure XP Beads (Beckman Coulter), pooled (384-plex) and 150bp paired-end reads were generated on a NovaSeq6000 (Illumina).

**Data analysis**

Fastq files were pseudo aligned to the mouse genome using Kallisto in Google Colabs (25). The matrix.abundance.gene.tpm.mtx file was used for further analysis and clustering using Seurat in R (26). Only nuclei with a number of unique genes greater than 5000 and a percentage of mitochondrial genes less than 0.5% were included in the clustering. Data were normalised using the *NormalizeData* command. Clustering in Seurat was performed based on 15 principal components and a resolution of 1.0 using the *FindClusters* command. Marker genes for each cluster were defined using *FindAllMarkers* with the minimum difference in the fraction of detection set to 0.1. Dotplots illustrating the expression of genes of interest across clusters were generated by cross referencing the 2000 most highly variable genes in our dataset (as identified by the *FindVariableFeatures* function) with lists of pain genes, neuropeptides, ion channels and GPCRs. List of pain genes were taken from the Pain Genes Database (62), Neuropeptides from neuropeptides.nl (63) and Ion channels and GPCRS from the IUPHAR database (64).

To compare transcriptomes between Naïve and SNI animals, nuclei were subsetted by mouse in order to generate counts for pseudobulk analysis. The

Deseq2 package in R was used to perform differential expression, considering genes with an adjusted p-value of less than 0.05.

Integration with other single cell/nucleus datasets was performed using Seurat (65). Data from Häring et al. was downloaded as molecule counts from GEO, and the excitatory clusters subsetted prior to integration and clustering. Data from Russ et al. were provided as a Robj Seurat Object by Ariel Levine. Again, excitatory neuron clusters were used for integration and clustering. Integration of the datasets was conducted as described in the Seurat vignette (66) with the addition of a regression command to attempt to account for the difference in sequencing depths between our dataset and others *(ScaleData(merged.integrated, verbose = FALSE, vars.to.regress = c("nFeature_RNA", "nCount_RNA"))).*

### RNAscope *in situ* hybridisation for spatial validation of clusters

Fluorescent *in situ* hybridisation was carried out with RNAscope probes and RNAscope Multiplex Fluorescent Assay v2 (Biotechne). Fresh frozen lumbar spinal cords from 4 Phox2a::Cre mice (two females, two males) were embedded in OCT medium and cut into 12 µm thick transverse sections with a cryostat (Leica CM1950). Sections were mounted non-sequentially (so that sections on the same slide were at least 4 apart) onto SuperFrost Plus slides (VWR) and air dried. Twelve sections from each animal were used for each of the three 3-plex reactions used for spatial validation (*Nmu/Cdh12/TdTomato*, *Baiap3/Cdh12/TdTomato* and *Gpr88/Erbb4/TdTomato)*. Reactions were performed according to the manufacturer's recommended protocol with several minor refinements. Slides were fixed for 1 h in 4% PFA and RNAscope Protease IV was applied for 15 minutes prior to hybridisation. Hybridisation products were visualised using TSA Vivid fluorophores (Tocris). TSA Vivid 520 was used for the C1 probes at a concentration of 1:1500. TSA Vivid 570 was used for the *TdTomato* probe at a concentration of 1:25 K. TSA Vivid 650 was used for the C3 probes at a concentration of 1:15 K Sections were mounted with Prolong-Glass anti-fade medium with NucBlue (Hoechst 33342) (ThermoFisher), stored at -20, and scanned with a Zeiss LSM710 confocal microscope equipped with Argon multi-line, 405 nm diode, 561 nm solid state and 633 nm HeNe lasers. Low magnification confocal scans of the TdTomato channel were taken of any sections containing Phox2a ALS cells, then higher magnification scans were obtained with a 40x (NA 1.3) oil-immersion lens of all 3 ISH fluorescence channels plus the nuclear staining. The laminar locations of the Phox2a cells were first plotted onto outlines of the dorsal horn using Neurolucida for Confocal software (MBF Bioscience). The transcriptional identity of each Phox2a cell was then recorded on these plots. Cells were only included if *TdTomato* transcripts were associated with a visible nucleus, and a cell was considered positive for expression of a gene if it contained >4 transcripts of mRNA. The individual plots were morphed to a standard outline of the dorsal horn in Inkscape (Inkscape Developers), resulting in single scalable vector graphics files for each probe combination in each animal. The overall percentages of neurons corresponding to ALS1-5 were normalized such that they added up to 100% for presentation in the main text and raw percentages are given in Figure S8.

**Combined IHC/ISH**

Combined immunohistochemistry/*in situ* hybridisation was conducted using perfusion fixed tissue from either Phox2a::Cre;Ai9 mice, or from Trpm8$^{Flp}$;RCE:FRT mice that had received injections of AAV.mCherry into the LPb. In all cases the animals were deeply anaesthetised and perfused with fixative, containing 4% freshly depolymerised formaldehyde in RNAase free buffers. Spinal cord segments were embedded in OCT and cut in either sagittal (Phox2a::Cre;Ai9) or horizontal section (Trpm8$^{Flp}$;RCE:FRT) on a cryostat (Leica CM1950). *In situ* hybridisation was conducted on slide using a RNAscope multiplex fluorescent kit according to manufacturer instructions with probes against *Nmu* or *Hs3st1.* Slides were not fixed further, and RNAscope Protease III was applied for 15 minutes prior to hybridisation. Hybridisation products were visualised using TSA Vivid 650 at a concentration of 1:3 K. Following *in situ* hybridisation, slides were rinsed 3 times in PBS, for ten minutes per rinse, and incubated overnight at RT (room temperature) in primary antibodies in PBS containing 0.3% Triton X-100 and 5% normal donkey serum (antibodies used in this study are detailed in table S2). Slides were then rinsed in PBS three times and incubated with secondary antibodies for 4 hours at RT. Secondary antibodies (Jackson Immunoresearch) were raised in donkey and conjugated to Alexa488 (concentration 1:500), and Rhodamine Red (concentration 1:100). Slides were rinsed again (3 x 10 minutes in PBS) prior to mounting with Prolong-Glass anti-fade medium (ThermoFisher) and storage at -20 ˚C. Sections were scanned with a Zeiss LSM900 Airyscan confocal microscope with 405, 488, 561 and 640 nm diode lasers. Confocal image stacks were obtained through a 40 × (NA 1.3) oil-immersion lens. All analyses were performed with Neurolucida for Confocal software (MBF Bioscience).

**Immunohistochemistry**

Multiple-labelling immunofluorescence reactions for the analysis of retrogradely labelled cells, and of the distribution of somatostatin and TRPM8-GFP were performed as described previously (19) on 60 μm thick transverse or horizontal sections of spinal cord, cut with vibrating blade microtomes (Leica VT1200 or VT1000). The sources and concentrations of antibodies are listed in Table S2.

Sections were incubated for 3 days at 4 ˚C in mixtures of primary antibodies diluted in PBS that contained 0.3 M NaCl, 0.3% Triton X-100 and 5% normal donkey serum, and then overnight in species-specific secondary antibodies (Jackson Immunoresearch, West Grove, PA, USA & Thermofisher for Alexa 'Plus' versions), which were raised in donkey and conjugated to Alexa488, Alexa488+, Alexa647, Rhodamine Red or Alexa405+. All secondary antibodies were diluted 1:500 in the same diluent, apart from those conjugated to Rhodamine Red which were diluted 1:100. Following immunoreaction, sections were mounted in anti-fade medium and stored at -20 °C. They were scanned with either a Zeiss LSM710 or LSM900 confocal microscopes.

For the analysis to test whether ALS cells in different laminae were immunoreactive for somatostatin, we first identified CTb-labelled cells in transverse sections of lumbar spinal cord and then determined whether these were positive for TdTomato (i.e. Phox2a-lineage cells). These classes (CTb+/Phox2a- and CTb+/Phox2a+) were plotted on outlines of the cord and their laminar locations documented. Antenna cells were identified by their characteristic pattern of dense peptidergic innervation using an antibody against substance P. The cells were then examined for somatostatin immunoreactivity.

For the analysis of the percentage of excitatory synaptic input to lamina I ALS cells arising from TRPM8-expressing afferents, we examined horizontal sections through lamina I from two $Trpm8^{Flp}$;RCE:FRT mice that had received LPb injections of AAV9.CB7.CI.mCherry.WPRE.rBG. Horizontal sections through lamina I were immunostained for mCherry, GFP and the postsynaptic density protein Homer (68). We assigned all of the retrogradely labelled cells to one of two classes: those with dense innervation from TRPM8 afferents, and those without. We then selected 10 lamina I ALS neurons from each animal, 5 from each class. Importantly, this selection was made by an investigator who was unaware of the distribution of GFP-labelling within the section. This was done to avoid bias towards cells with extreme patterns of input. The dendritic arbors of the cells were then reconstructed by a separate investigator. We used Homer immunoreactivity to identify excitatory synapses onto these cells. We then determined the percentage of Homer puncta on each cell that were adjacent to a TRPM8 (GFP-positive) primary afferent bouton. All analyses were performed with Neurolucida for Confocal software (MBF Bioscience).

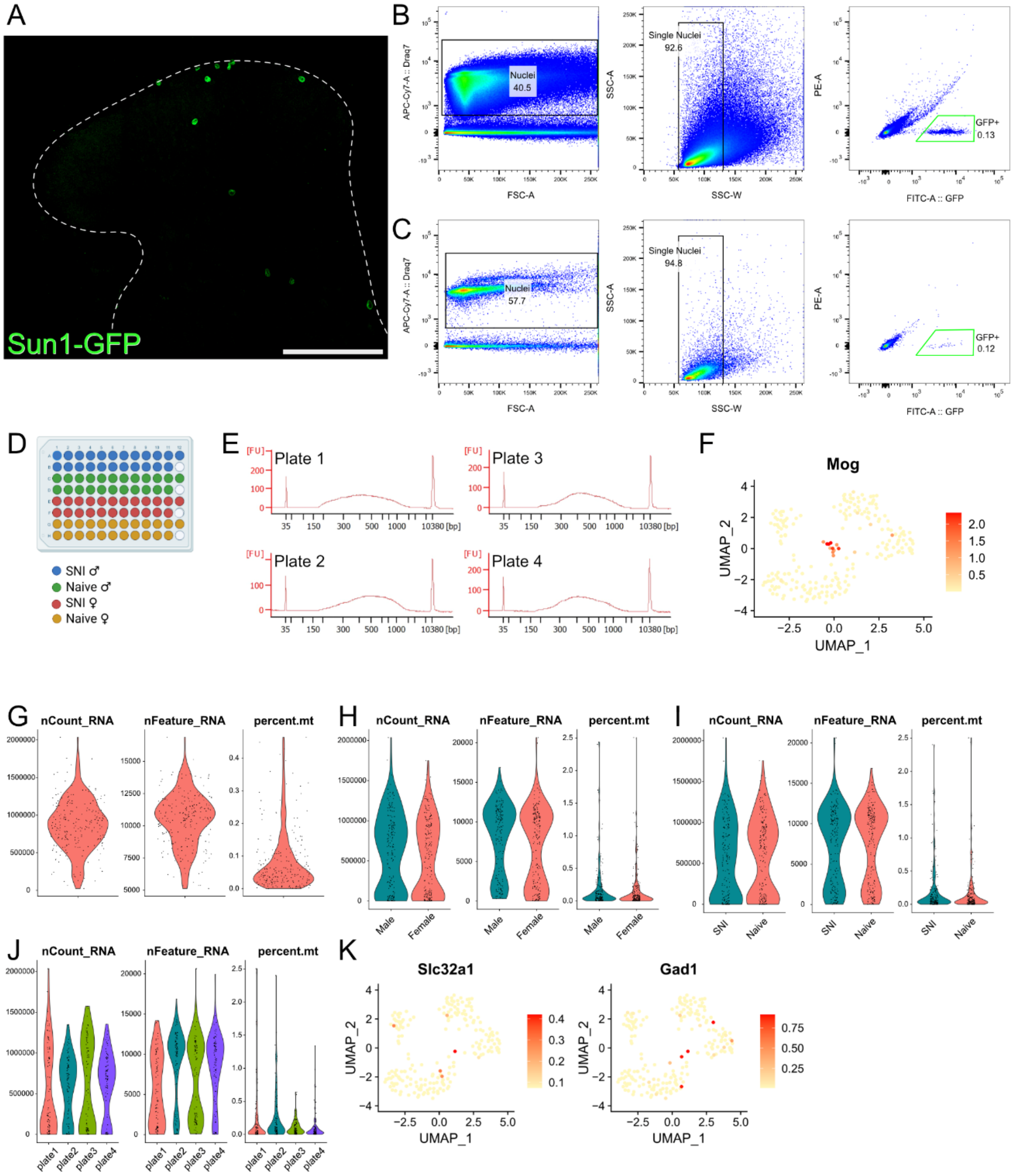

**Fig. S1. Data to support the sequencing approach used.** (**A**) A transverse image of the L4 segment of a Phox2a::Cre;Sun1-GFP mouse. The section has been immunoreacted for GFP and shows several Phox2a nuclei in both the superficial and deep dorsal horn. The image is a projection of a confocal image stack taken through an entire 60µm thick section at 2µm z-spacing. Scale bar = 100 µm. (**B** & **C**) Gating strategy used for fluorescence-activated cell sorting (FACS) of Phox2a-GFP nuclei. B shows concatenated data from all 16 animals in the study**.** C shows the original data from a single representative animal. (**D**) To avoid batch effects, we used a balanced plate layout, exemplified here, for each of the 4 plates submitted for Smart-seq2 RNA sequencing. Each well contained a single nucleus. (**E**) Bioanalyzer traces from each plate taken to assess cDNA quality prior to pooling of the 4 libraries prior to sequencing. (**F**) A small cluster of nuclei defined by expression of the oligodendrocyte marker *Mog.* These nuclei were removed prior to analysis. (**G**–**J**) Violin plots showing total number of genes (nCount_RNA), number of unique genes (nFeature_RNA) and percentage mitochondrial genes (percent.mt) detected per nucleus. G shows these metrics for the 206 nuclei submitted to the final analysis after QC. H-J show these data for all nuclei sequenced prior to QC split by sex, condition and plate. (**K**) UMAP feature plots demonstrating minimal expression of two canonical markers of inhibitory neurons in the 206 nuclei in the analysis.

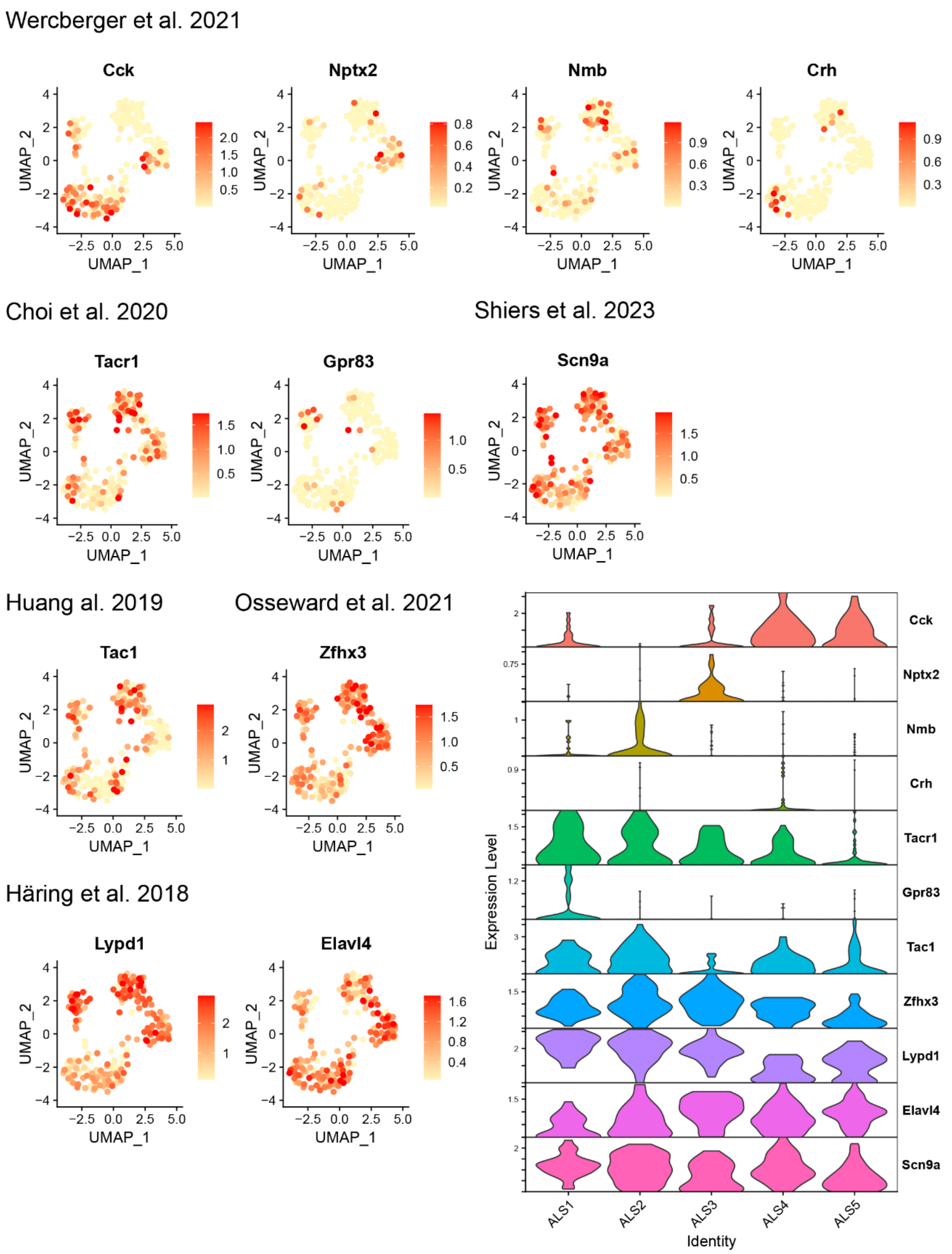

**Fig. S2. Distributions of existing projection neuron markers.** The expression of marker genes for ALS neurons and subsets of ALS neurons proposed in previous studies. For each gene, a UMAP feature plot shows the expression of mRNAs across the 206 nuclei in this study. Violin plots also show the expression of each mRNA across ALS 1-5. All studies are cited within the main text of the article except Shiers et al. 2023, which is cited below (67).

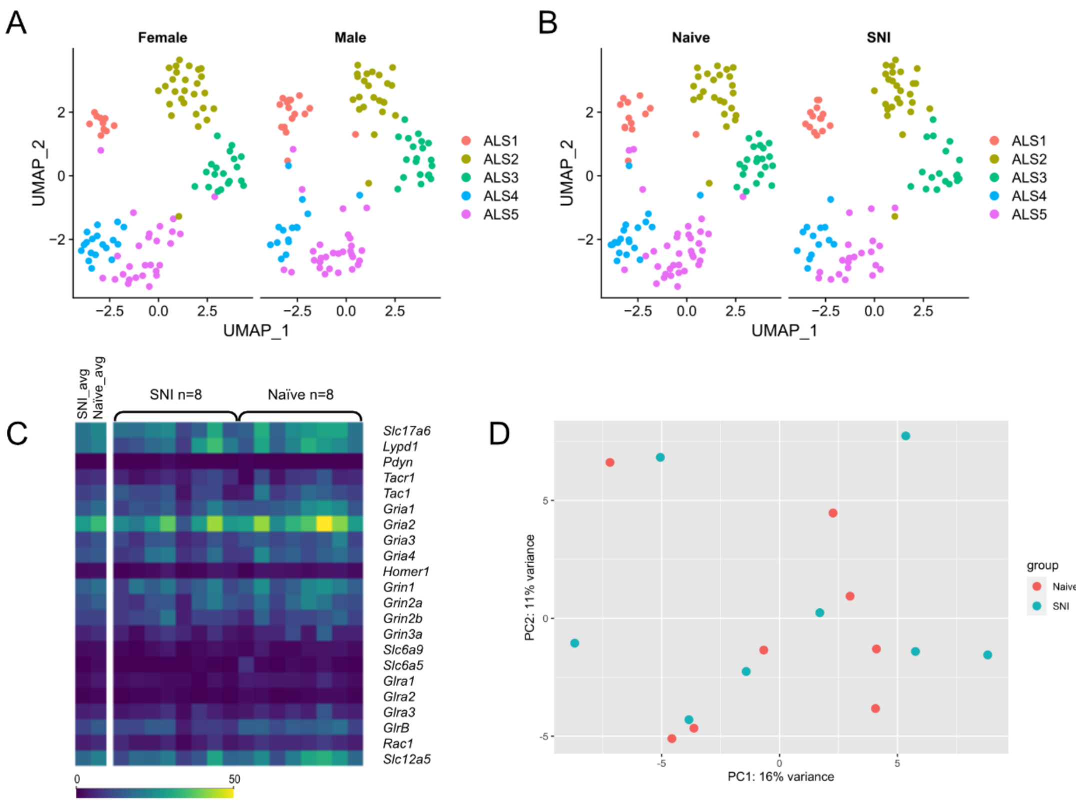

**Fig. S3. Lack of evidence that SNI alters the ALS transcriptome.** (**A**) UMAP plots showing the distribution of ALS nuclei into clusters split by sex. (**B**) UMAP plots showing the distribution of ALS nuclei into clusters split by condition (SNI/Naïve). (**C**) Heatmap showing pseudobulk normalized relative counts across animals and conditions for selected genes with possible roles in projection neuron identity and signaling. No significant differences were found using differential expression analysis across the whole transcriptome. (**D**) PCA plot showing principal components based on pseudobulk gene expression for SNI and Naïve animals (n = 8 per group).

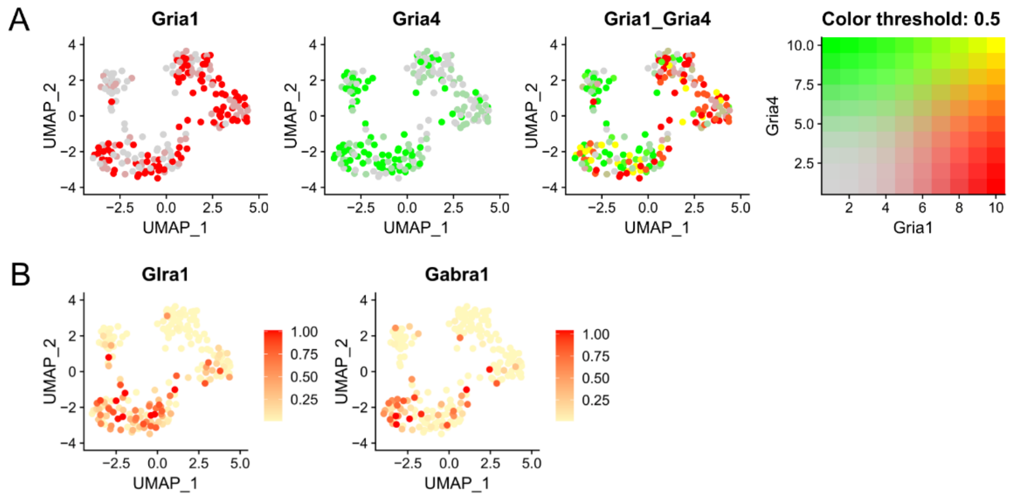

**Fig. S4 Expression of ionotropic receptor subunits across clusters** (A) UMAP feature plots showing the expression of mRNAs for *Gria1* and *Gria4* across ALS clusters. Cells are coloured by the expression levels of each mRNA in the first two panels, then by the expression level of both in the third. (B) UMAP feature plots showing the expression of *Glra1* and *Gabra1*.

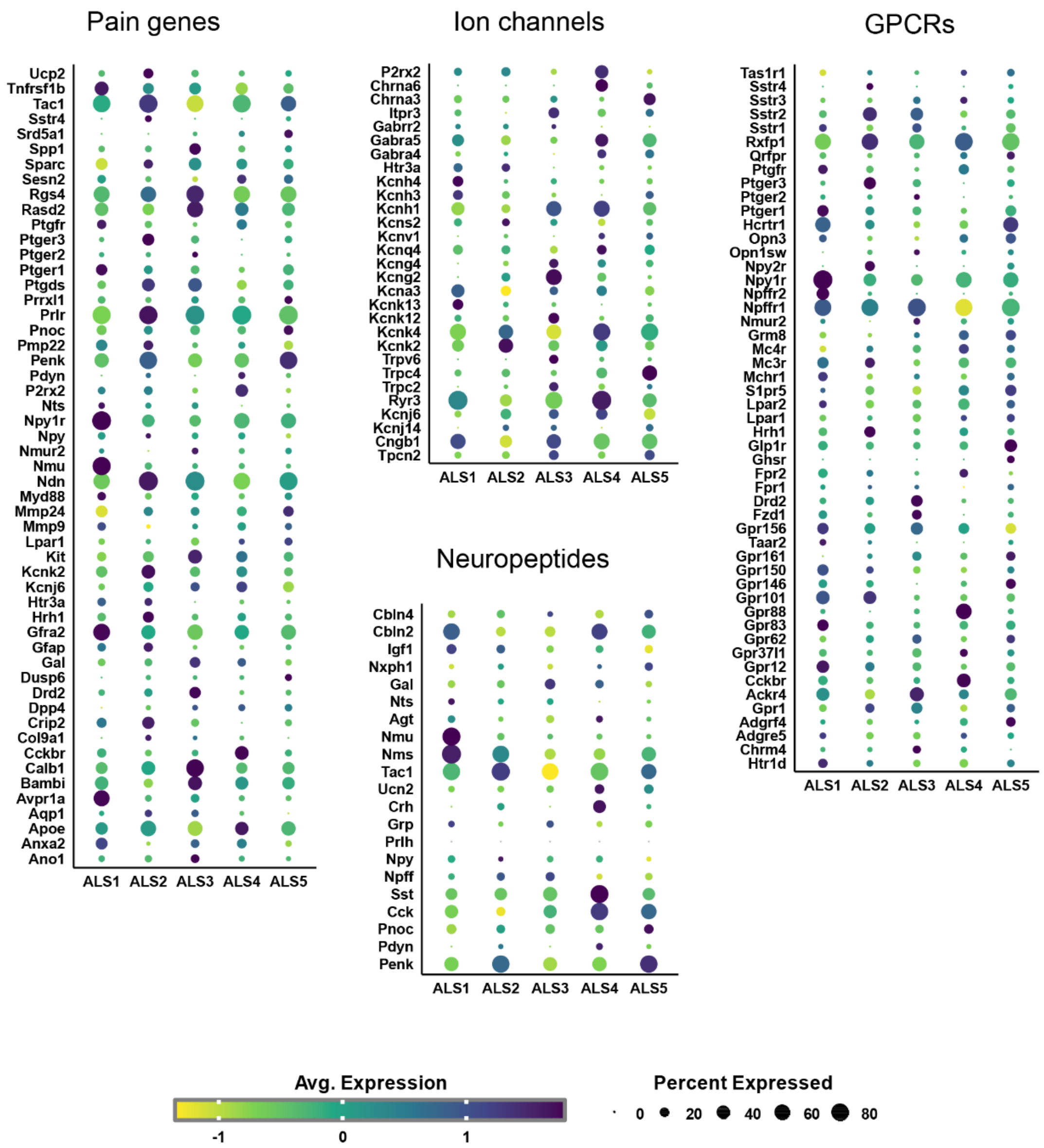

**Fig. S5. Expression of select functional classes of genes in each cluster.** Dot plots showing the expression of genes across ALS1-5. The colour of each dot represents the z-score of the scaled gene expression within the cluster and the size of each dot represents the percentage of cells in each cluster expressing the listed gene. Genes were chosen by cross referencing the 2000 most variable genes in our dataset with databases of pain genes, neuropeptides, voltage and ligand-gated ion channels and G-protein linked receptors.

### A - Integration with Häring et al.

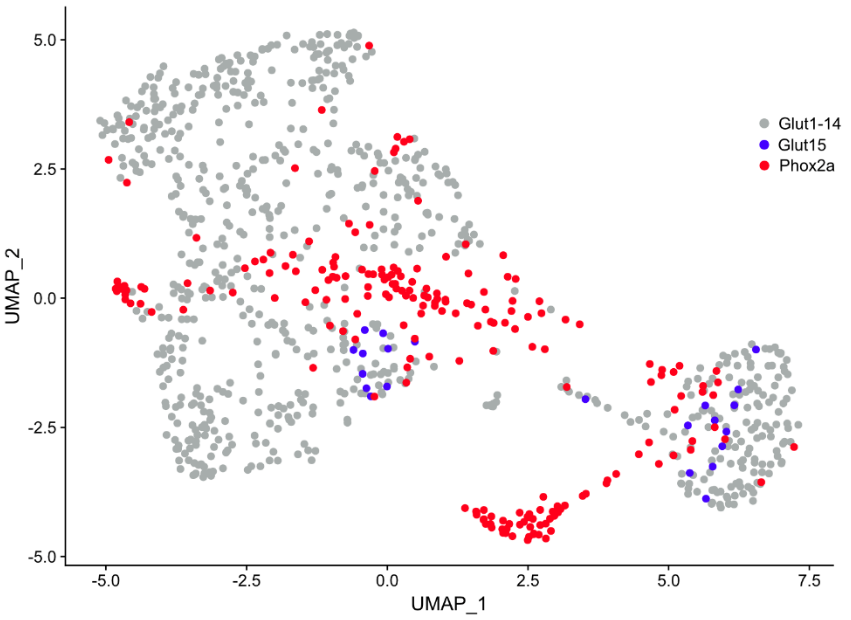

### B - Integration with Russ et al.

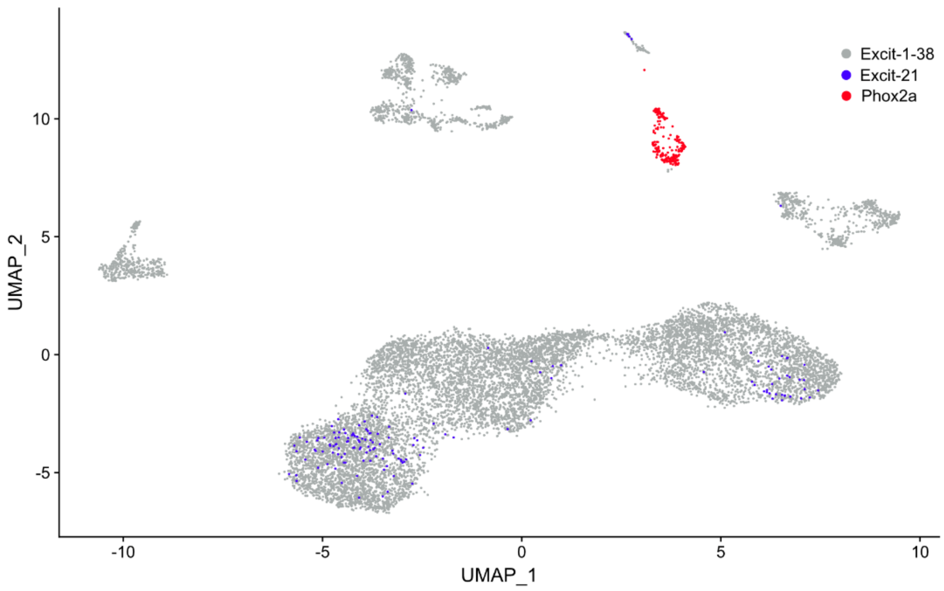

**Fig. S6. Integration of 206 Phox2a nuclei with existing spinal cord single-cell/nucleus datasets.** UMAP plots showing the integration of the dataset generated here with the excitatory neuron subsets of two dorsal horn transcriptomic atlases using Seurat. In each case, the Phox2a nuclei are shown in red, the excitatory neurons from the existing atlases in grey and the putative projection neuron cells from the atlases in blue.

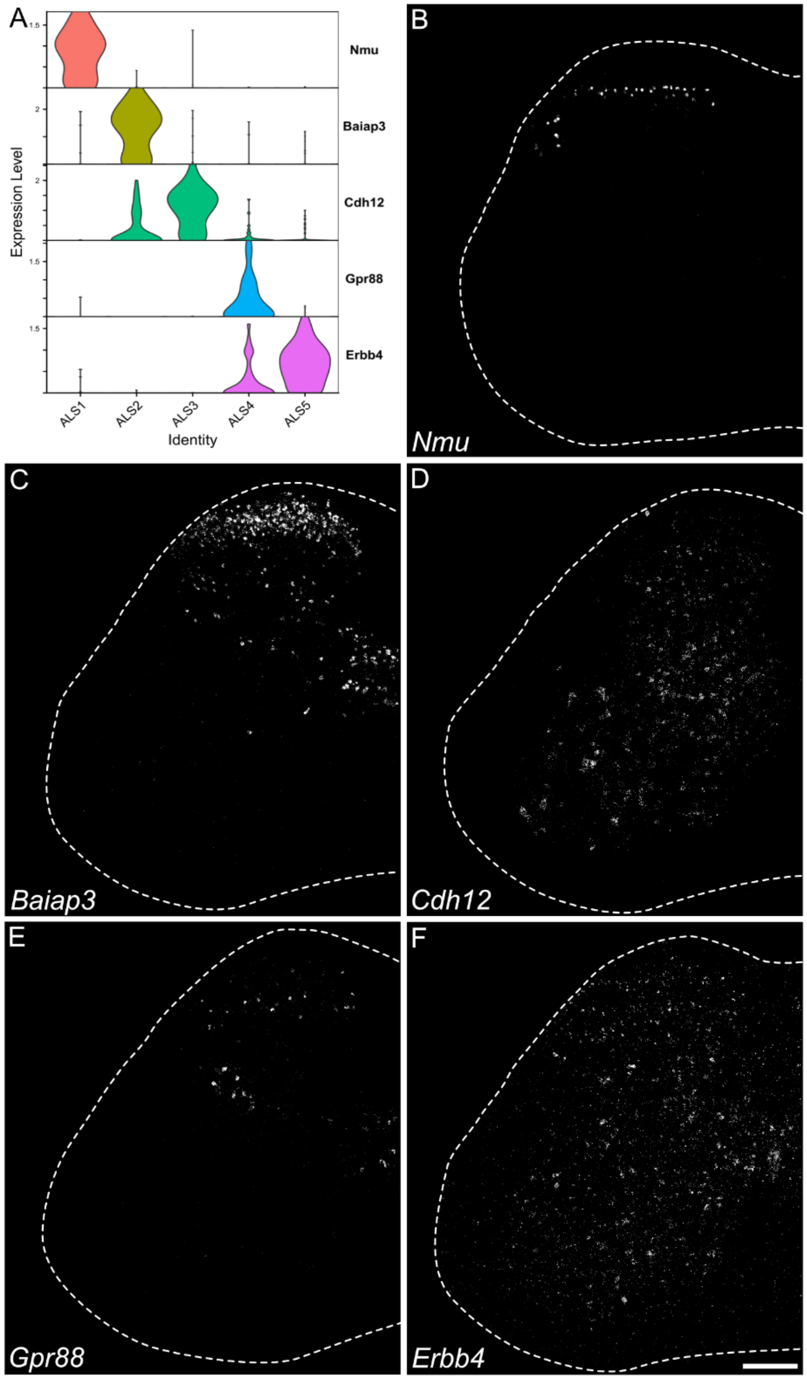

**Fig. S7. Marker genes used for *in situ* validation of Phox2a clusters.** (**A**) Violin plots showing the expression of the five markers used for *in situ* validation experiments in each cluster. (**B**-**F**) Low power confocal images of RNAscope *in situ* hybridization showing the distribution of mRNAs for each of the markers across transverse sections of the lumbar spinal cord. In each case, the images are projections of confocal z-stacks through the entire 12 μm section (2 μm z-spacing). The outline indicates the edge of the white matter. Scale bar = 200 μm.

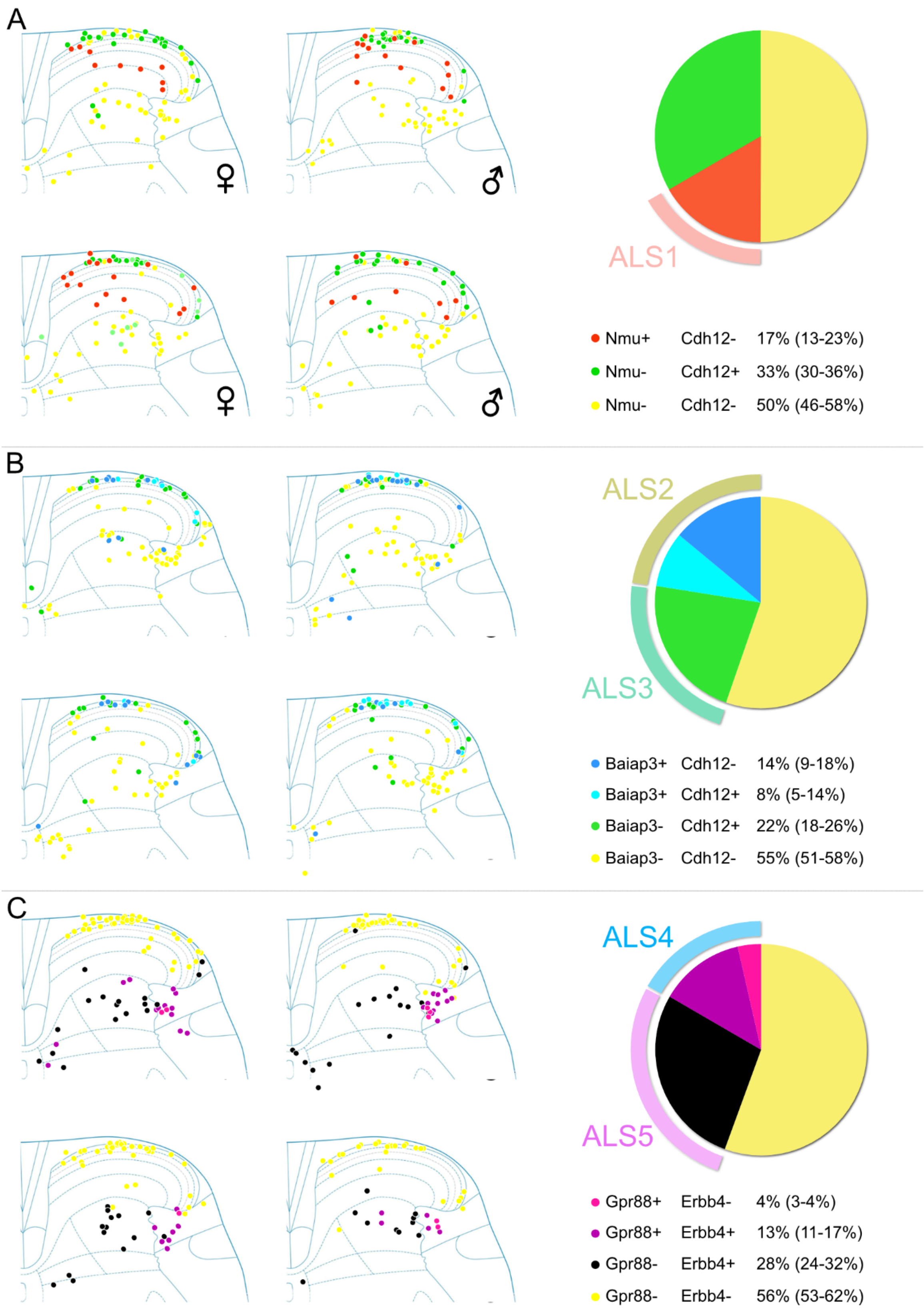
A
♀
♂
ALS1
Nmu+ Cdh12- 17% (13-23%)
Nmu- Cdh12+ 33% (30-36%)
Nmu- Cdh12- 50% (46-58%)
B
ALS2
ALS3
Baiap3+ Cdh12- 14% (9-18%)
Baiap3+ Cdh12+ 8% (5-14%)
Baiap3- Cdh12+ 22% (18-26%)
Baiap3- Cdh12- 55% (51-58%)
C
ALS4
ALS5
Gpr88+ Erbb4- 4% (3-4%)
Gpr88+ Erbb4+ 13% (11-17%)
Gpr88- Erbb4+ 28% (24-32%)
Gpr88- Erbb4- 56% (53-62%)

**Fig. S8. *In situ* validation of Phox2a ALS clusters.** Each section (**A**-**C**) shows the distribution and overlap of marker gene expression for the three pairs of markers used for *in situ* hybridization spatial validation experiments. Each dot represents a Phox2a TdTomato+ cell and separate plots are shown representing each of the 4 animals (of both sexes) that were used in the analysis. The colour of the dot represents the classification of the cell according to the marker gene expression, and cells negative for both genes are shown in yellow. The pie charts show the average proportion (and range in brackets) for each class of cells across the 4 animals with the cluster allocation indicated by the detached segments at the circumference. Note, the percentages given here for each cluster are slightly different to those given in the main text. In the main text, the percentages corresponding to ALS1-5 have been normalized such that they add up to 100%. The sum of percentages here is 106% as these were obtained from different tissue sections reacted with different probes. This normalization resulted in the following percentage of cells in each cluster; ALS1 16%, ALS2 21%, ALS3 21%, ALS4 16%, ALS5 26%.

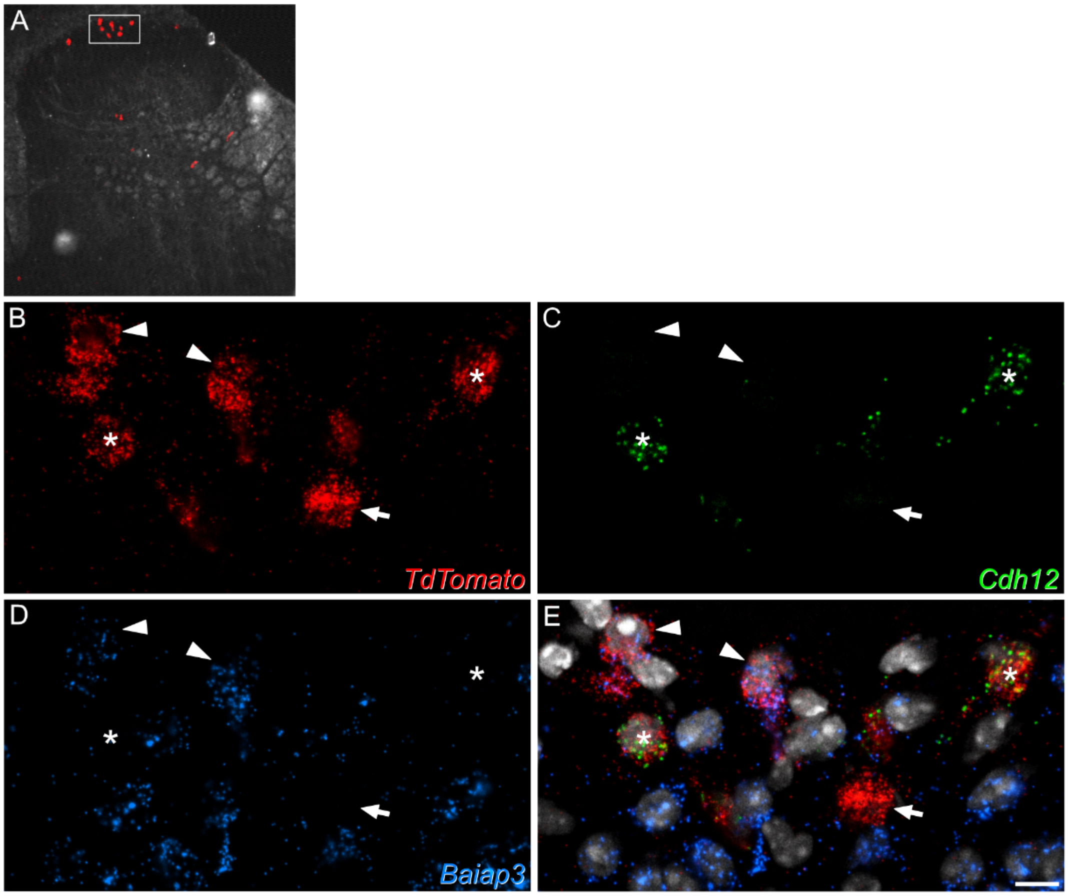

**Fig. S9. A rare cluster of lamina I cells illustrates the ALS2 and ALS3 populations.** (**A**) A low magnification confocal image of RNAscope *in situ* hybridization. mRNAs for Tdtomato are shown combined with a dark field image of a transverse lumbar spinal cord section from a Phox2a::Cre;Ai9 mouse. The box highlights an area where several lamina I Phox2a ALS neurons are present. This area is enlarged in high magnification scans in panels **B**-**E.** The mRNAs for *Cdh12* and *Baiap3* have been revealed alongside the *TdTomato* mRNA, and these are shown combined in E**.** Five Phox2a+ ALS neurons can be seen. Two (marked by arrowheads) are positive for *Baiap3* and negative for *Cdh12*, and therefore belong to ALS2. Two (marked with asterisks) are positive for *Cdh12* and negative for Baiap3 and belong to *ALS3*. A single cell (marked with an arrow) is negative for both transcripts and could represent an ALS1 cell. Image is a projection of a confocal z-stack taken through the entire depth of a 12μm section (1 μm z-spacing). Scale bar = 10 μm.

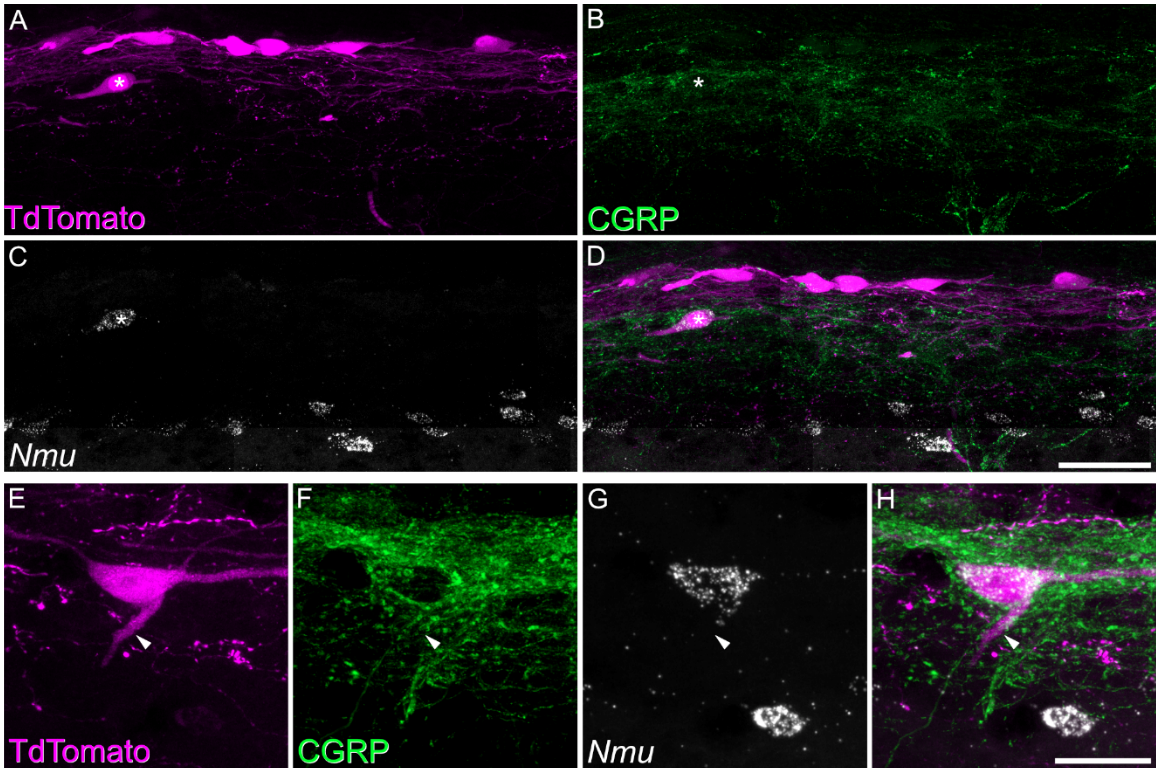

**Fig. S10. Distinct characteristics of ALS1 cells in lamina I.** (**A**-**D**) Combined *in situ* hybridisation and immunohistochemistry performed on a sagittal section of lumbar spinal cord from a Phox2a::Cre;Ai9 mouse. A and B show immunostaining for TdTomato and CGRP, while C shows *in situ* hybridisation signal for *Nmu*, and D is a merged image. A large TdTomato labelled lamina I cell is indicated with an asterisk. *In situ* hybridisation shows that this cell possesses *Nmu* mRNA and therefore belongs to the ALS1 cluster. The ALS1 cell illustrated is adjacent to several other TdTomato labelled ALS neurons in lamina I which are negative for *Nmu*. The ALS1 cell is characteristically positioned deeper in lamina I than the other cells. (**E**-**H**) Images show the same combined *in situ* hybridisation and immunohistochemistry technique applied to a different tissue section. Another lamina I Phox2a neuron is shown, and again this cell expresses *Nmu* and belongs to ALS1. In this case, the ventrally directed dendrite of the cell (arrowhead) is associated with peptidergic CGRP bundles. In both cases, images are a projection of confocal optical sections (1 μm z-separation) taken through the full 12 μm thickness of the section. Scale bar = 50 μm in A-D and 10 μm in E-H.

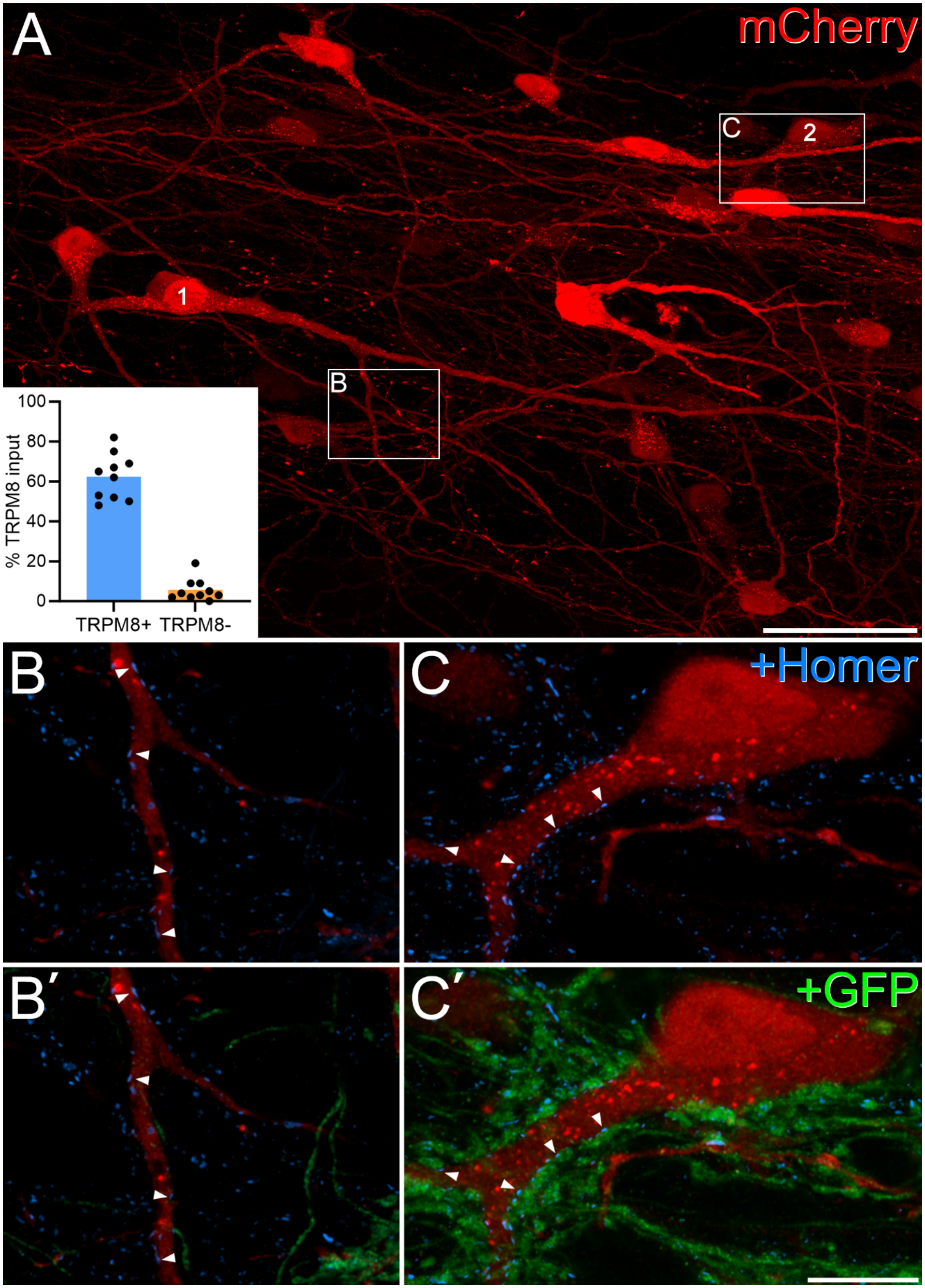

**Fig S11. Quantification of synaptic input to lamina I ALS neurons arising from TRPM8-expressing afferents. A**: A horizontal section through lamina I from a Trpm8$^{Flp}$;RCE:FRT mouse that had received an injection of AAV.mCherry into the contralateral lateral parabrachial area. Several retrogradely labelled cells are visible, and two of these are marked (1, 2). Boxes indicate areas seen at higher magnification in **B** and **C**. The inset shows quantification of the percentage of excitatory synapses (revealed by the presence of Homer) that were associated with TRPM8-positive (GFP-labelled) boutons for 10 cells classified as densely innervated by TRPM8 afferents (blue bar, TRPM8+) and 10 cells classified as not densely innervated by TRPM8 afferents (orange bar, TRPM8-). **B**, **B'**: Higher magnification views of the regions marked by the boxes in **A**, scanned to reveal Homer (blue). **B** shows part of the dendritic tree of cell 1, while **C** includes the cell body and proximal dendrite of cell 2. Several Homer puncta (corresponding to postsynaptic densities of excitatory synapses) can be seen on each of the cells, and some of these are marked with arrowheads. **B'**, **C'**: The same fields showing both immunoreactivity for both Homer (blue) and GFP (green). All of the marked synapses on cell 2 are associated with GFP-labelled (TRPM8-positive) boutons, whereas none of the marked synapses on cell 1 are contacted by GFP-labelled boutons. Image are maximum intensity projection of 103 (**A**), 3 (**B**, **B'**) or 4 (**C**, **C'**) optical sections at 0.5 μm z-spacing. Scale bars = 50 μm (**A**) and 10 μm (**B**, **C**).

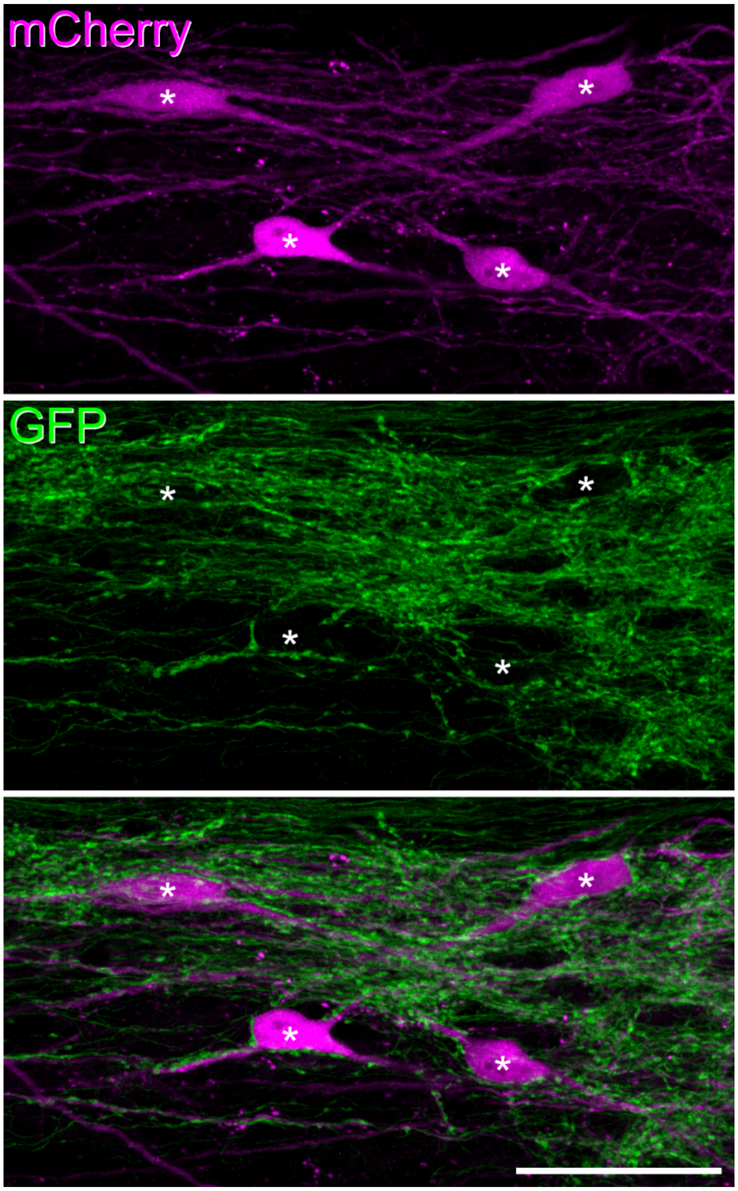

**Fig. S12. Phox2a-lineage cells that are associated with TRPM8 afferents.** A horizontal section through lamina I from a Phox2a::Cre;Ai9;Trpm8$^{Flp}$;RCE:FRT mouse, in which Phox2a+ ALS neurons are labelled with mCherry and TRPM8 afferents with GFP. Four labelled lamina I cells are present in this field (marked with asterisks), and these lie within a bundle of TRPM8 afferents. All four of these cells are contacted by numerous TRPM8-positive boutons. The images are maximum intensity projections of 7 confocal optical sections at 0.5 µm z-spacing. Scale bar = 50 µm.

**Table S1.** The proportion of CTb-labelled and Phox2a-positive ALS projection neurons that were somatostatin-immunoreactive.

| Region | No. CTb+ | No. Phox2a+ | %Sst/CTb | %Sst/Phox2a |
|---|---|---|---|---|
| Lamina I | 91<br>(79-103) | 50<br>(47-53) | 8.5%<br>(6.3-10%) | 6.1%<br>(2.2-10%) |
| Antenna | 9<br>(4-14) | 9.5<br>(5-14) | 0.0%<br>(0-0%) | 0.0%<br>(0-0%) |
| Lateral lamina V | 86.5<br>(60-113) | 19<br>(11-27) | 54.1%<br>(53-55%) | 48.1%<br>(46-50%) |
| Medial lamina V | 16<br>(13-19) | 11.5<br>(7-16) | 6.5%<br>(5.3-7.7%) | 4.2%<br>(0-8.3%) |
| Central canal | 67<br>(57-77) | 11<br>(6-16) | 1.5%<br>(1.3-1.8%) | 3.8%<br>(0-7.7%) |
| Lateral spinal nucleus | 34.5<br>(25-44) | 5.5<br>(1-9) | 24.8%<br>(20-29%) | 55.6%<br>(11-100%)* |

Data were obtained from 2 Phox2a::Cre;Ai9 mice in which CTb had been injected into the lateral parabrachial area. *The 100% value for the lateral spinal nucleus was from a mouse in which only 1 Phox2a-positive cell was present in this region. The percentage of CTb labelled cells expressing somatostatin for projection neurons in all areas except lateral lamina V was 9%.

**Table S2.** Antibodies used in this study.

| Antigen | Host species | Source | Cat no | Dilution | RRID | Expt type |
|---|---|---|---|---|---|---|
| Somatostatin | Rabbit | Peninsula | T-4103 | 1:1000 | RRID:AB_518614 | Somatostatin quantification |
| CTb | Goat | List Biological | 703 | 1:5000 | RRID:AB_10013220 | Somatostatin quantification |
| mCherry | Chicken | Abcam | Ab205402 | 1:10,000 | RRID:AB_2722769 | Somatostatin quantification |
| Substance P | Rat | Oxford biotechnology | OBT06435 | 1:200 | N/A | Somatostatin quantification |
| mCherry | Rat | Invitrogen | M11217 | 1:1000 | RRID:AB_2536611 | Combined IHC/FISH |
| GFP | Chicken | Abcam | ab13970 | 1:1000 | RRID:AB_300798 | Combined IHC/FISH |
| CGRP | Guinea pig | Bachem | T-5027 | 1:1000 | RRID:AB_518152 | Combined IHC/FISH |
| GFP | Rabbit | Watanabe | | 1:1000 | RRID:AB_2491093 | TRPM8/GFP analysis |
| Homer1 | Goat | Watanabe | | 1:1000 | RRID:AB_2631104 | TRPM8/GFP analysis |

**Table S3.** Key resources table.

| Reagent or Resource | Source | Identifier |
|---|---|---|
| **Mouse Lines** | | |
| **Ai9(RCL-tDT)** | Jax | Strain 007909<br>RRID:IMSR_JAX:007909 |
| **CAG-Sun1/sfGFP known as Sun1-GFP** | Jax | Strain 021039<br>RRID:IMSR_JAX:021039 |
| **RCE:FRT** | Jax | Strain 032038-JAX<br>RRID:MMRRC_032038-JAX |
| **Phox2a::Cre** | Artur Kania | N/A |
| **Sst-IRES-Cre - known as $Sst^{Cre}$** | Jax | Strain 013044<br>RRID:IMSR_JAX:013044 |
| **Cck-IRES-Cre - known as $Cck^{Cre}$** | Jax | Strain 012706<br>RRID:IMSR_JAX:012706 |
| **$Trpm8^{Flp}$** | Mark Hoon | N/A |
| **Reagents and labware** | | |
| **DRAQ7 Dye** | Invitrogen | D15105 |
| **RNAsin** | Promega | N2111 |
| **"cOmplete™, Mini, EDTA-free Protease Inhibitor Cocktail** | Roche | 11836170001 |
| **Twin.tec 96 Well LoBind PCR Plates, Skirted** | Eppendorf | 15280735 |
| **NextEra XT kit** | Illumina | FC-131-1096 |
| **AMPure XP Beads** | Beckman Coulter | A63881 |
| **Superfrost Plus slides** | VWR | 48,311–703 |
| ***In Situ* Hybridisation** | | |
| **RNAscope Multiplex Fluorescent Assay v2** | Biotechne | 323100 |
| **TSA Vivid fluorophores 520/570/650** | Tocris | 7523/7526/7527 |
| **RNAscope Probe - Mm-Baiap3-C3** | Biotechne | 1133511-C3 |
| **RNAscope Probe - Mm-Cdh12** | Biotechne | 842531 |
| **RNAscope Probe - Mm-Erbb4-C3** | Biotechne | 318721-C3 |
| **RNAscope Probe - Mm-Nmu-C3** | Biotechne | 446831-C3 |
| **RNAscope Probe - Mm Gpr83** | Biotechne | 317451 |
| **RNAscope Probe - TdTomato** | Biotechne | 317041-C2 |
| **RNAscope Probe - Mm-Hs3st1-C2** | Biotechne | 570431-C2 |

| | | |
|---|---|---|
| **Equipment** | | |
| **Bioanalyzer** | Agilent | 5067-4626 |
| **Mosquito LV** | SPT Labtech | N/A |
| **Biomek NX Automated WorkStation** | Beckman Coulter | N/A |
| **Leica CM1950 cryostat** | Leica | N/A |
| **Viruses** | | |
| **AAV9.CB7.CI.mCherry.WPRE.rBG - Known as AAV9.mCherry** | Penn Core | V3874TI-R |
| **ssAAV-1/2-CAG-dlox-tdTomato(rev)-dlox-WPRE-bGHp(A) - Known as AAV9.flex.TdTomato** | VVF-Zurich | v167-9 |
| **Software** | | |
| **R 4.2.2** | The R Project | |
| **Kallisto** | Pacter Lab | |
| **Seurat v4.1.1** | Satija Lab | |
| **Neurolucida 2021.1.3** | MBF Bioscience | |
| **Zen 3.4** | Zeiss microscopy | |
| **Inkscape 1.2** | Inkscape developers | |
